# Rbpms2 promotes female fate upstream of the nutrient sensing Gator2 complex component, Mios

**DOI:** 10.1101/2024.01.25.577235

**Authors:** Miranda L. Wilson, Shannon N. Romano, Nitya Khatri, Devora Aharon, Yulong Liu, Odelya H. Kaufman, Bruce W. Draper, Florence L. Marlow

## Abstract

Reproductive success relies on proper establishment and maintenance of biological sex. In many animals, including mammals, the primary gonad is initially ovary in character. We previously showed the RNA binding protein (RNAbp), Rbpms2, is required for ovary fate in zebrafish. Here, we identified Rbpms2 targets in oocytes *(Rbpms2-bound oocyte RNAs; rboRNAs*). We identify Rbpms2 as a translational regulator of *rboRNAs*, which include testis factors and ribosome biogenesis factors. Further, genetic analyses indicate that Rbpms2 promotes nucleolar amplification via the mTorc1 signaling pathway, specifically through the mTorc1-activating Gap activity towards Rags 2 (Gator2) component, Missing oocyte (Mios). Cumulatively, our findings indicate that early gonocytes are in a dual poised, bipotential state in which Rbpms2 acts as a binary fate-switch. Specifically, Rbpms2 represses testis factors and promotes oocyte factors to promote oocyte progression through an essential Gator2-mediated checkpoint, thereby integrating regulation of sexual differentiation factors and nutritional availability pathways in zebrafish oogenesis.

## Introduction

Oogenesis is an essential process for all sexually reproducing organisms. Successful oogenesis is complex and demanding, requiring immense cellular growth, high transcriptional and translational burdens, and the generation of a large reserve of maternal factors necessary for the embryo. In some organisms, such as humans, oocytes are very long-lived cells that must tightly regulate their development and maturation in response to internal and external cues (reviewed in^1^). Our lab has previously shown that the RNA binding protein of multiple splice forms 2 (Rbpms2) is required for oocyte development and subsequent female sex differentiation in zebrafish but its mode of action and targets remain unclear^2,3^. Based on our prior findings that Rbpms2 protein is expressed in oocytes but not detected in somatic gonad or testis^2^, is required for ovary development, and promotes ovary fates independent of the checkpoint regulators Tp53 and Chk2^2^, we envisioned two non-mutually exclusive hypotheses for Rbpms2 functions in ovary specific differentiation of germ cells. 1) Rbpms2 actively represses translation of testis factors that are expressed in the early germline cells of ovaries and 2) promotes translation of factors that allow for oocyte growth and differentiation upstream of an unknown oocyte progression checkpoint.

Early oogenesis relies on a high degree of transcription and generation of translation machinery components, like ribosomes, to allow for extreme growth and support the translational needs of oocytes (reviewed in^4^). Therefore, adequate nutrition is necessary for oogenesis (reviewed in^5^). The interplay of oocyte development, nutrition availability, and fertility has been observed across species^6–8^, including zebrafish where a low nutrition environment biases sexual development to male^9^. In other species, low nutrition environments can suppress or arrest oogenesis to preserve energy for homeostatic mechanisms^6–8^.

Nutrition sensing pathways are important to the viability of all cells. For example, high levels of fatty acids and carbohydrates are necessary for activation of the citric acid cycle for adenosine triphosphate (ATP) production whereas glycolysis relies on sufficient glucose intake. Other mechanisms respond to sufficient amino acid levels, like the metabolic homeostasis regulator mechanistic target of rapamycin (mTOR) pathway (reviewed in^10^). Through amino acid detection, the Gap activity towards Rags 2 (GATOR2) complex inhibits the Gap activity towards Rags 1 (GATOR1), prompting activation of the mTOR complex 1 (mTORC1) which induces a signaling cascade that positively regulates cell growth, metabolism, and autophagy (reviewed in^10,11^). Conversely, metabolic stress signals like hypoxia prompt activation of the tuberous sclerosis 1 (TSC1) and 2 (TSC2) complex, which inhibits mTORC1 signaling by inactivation of Ras homolog, mTORC1 binding (Rheb) (reviewed in^10^). Notably, dysregulation of mTORC1 signaling in oocytes has been shown to have varying impacts on short- and long-term fertility across organisms through poorly understood mechanisms^12–17^. For example, in *Drosophila* ovarioles with reduced mTORC1 signaling due to knockout of the GATOR2 complex protein, Missing oocyte (*mio*; *mios* in zebrafish), oocytes are initially specified but not maintained^18^. How mTORC1 signaling contributes to sustained oogenesis remains unclear. In other cell types, mTORC1 signaling has been shown to support early stages of ribosome biogenesis^19^. Sufficient ribosome biogenesis in oocytes is vital to support their translational burden and to provide the ribosomes the embryo requires (reviewed in^4^). However, the interplay between nutrition, mTORC1 signaling, and ribosome biogenesis in the context of oocyte development remains to be fully understood.

Here we identified a role for the vertebrate-specific oocyte fate regulator Rbpms2 in nucleolar expansion and mTorc1 signaling. Using an oocyte specific tagged-Rbpms2 for RNA immunoprecipitation, we found that Rbpms2 target RNAs in oocytes *(Rbpms-bound oocyte RNAs; rboRNAs*) include regulators of RNA metabolism and degradation, regulators of ribosome biogenesis, and surprisingly, RNAs required for testis development. Cross-referencing the *rboRNAs* to previously published single cell RNA-sequencing ovary data^20^, we identified an enrichment of *rboRNAs* in early oocytes, the stages crucial for further oocyte and ovary differentiation in zebrafish. Subsequent RNA-seq analysis implicates Rbpms2 as a *rboRNA* translational regulator, likely acting to repress testis factors and activate ovary factors. Further, RNA-seq and immunohistological analyses indicate that Rbpms2 promotes nucleolar assembly, ribosome biogenesis, and mTorc1 pathway components. Thus, we hypothesized that early gonocytes are in a dual poised bipotential state where Rbpms2 dynamically regulates oocyte and testis *rboRNAs* and acts as a switch upstream of a nutrient responsive pathway, positively impacting the translational capacity of oocytes.

Accordingly, genetic analyses place Rbpms2 upstream of the mTorc1 pathway-activating protein, Mios. Furthermore, Mios loss, like loss of Rbpms2, disrupts nucleolar composition and compromises ovary maintenance. Genetically manipulating regulators of the metabolic stress sensing arm of mTorc1 in *mios* mutants (*mios^−/−^*) did not prevent oocyte loss, while active forms of mTOR restored oogenesis and female sex determination in *mios*^−/−^ fish. Cumulatively, we have identified an RNA binding protein (RNAbp)-mediated binary fate switch that activates a Gator2-mediated checkpoint essential for oocyte development, thereby integrating sexual differentiation factors and nutritional availability pathways during zebrafish oogenesis and female sex determination.

## Results

### Rbpms2 binds to factors required for testis development that are expressed in early oocytes

In nongonadal somatic cells, Rbpms family proteins are thought to act primarily as translational regulators. Recently this has been shown during cardiac commitment of human embryonic stem cells (hESCs) where RBPMS controls the translation initiation of several factors, and subsequent developmental pathways, by recruitment to actively translating ribosomes^21^. However, the germline-specific function(s) and RNA target(s) of Rbpms2 have yet to be characterized^22^. Therefore, we sought to determine the germline-specific RNA targets of Rbpms2, which are expected to be critical for successful oogenesis and reproductive functions.

As Rbpms2 is expressed in multiple tissues^2^, we took a transgenic RNA immunoprecipitation (RNAIP) approach to isolate Rbpms2 target RNAs in oocytes. Briefly, we used previously generated stable adult transgenic fish expressing mApple-Rbpms2 or mApple alone under the oocyte specific promoter, *buckyball* (Figure 1a)^2,23^. This promoter was selected because it is specifically expressed in oocytes but not ovarian somatic cells^23,24^. We identified 732 RNAs bound by Rbpms2 in oocytes of adult ovaries (*rboRNAs*) (Figure 1b). Among *rboRNAs,* 149 have been associated with spermatogenesis and testis fate (*rbtRNAs*) in zebrafish and other organisms (Figure 1b, Supp. Figure 1a). Mapping the temporal expression of *rboRNAs* to previously generated 40 days post fertilization (dpf) zebrafish ovary single cell RNA-sequencing data^20^ revealed that 680 of these RNAs are expressed throughout mitotic, meiotic, and early oocyte cells (Figure 1b). The remaining 52 *rboRNAs* that did not map to the 40 dpf ovary dataset are likely transcripts expressed in later stage oocytes present in the fully mature adult ovary. Notably, several *rbtRNAs* were enriched in the undifferentiated mitotic and pre-meiotic cells of the 40 dpf zebrafish ovary (Figure 1b)^20^. This is consistent with our previous findings that *rbpms2* DMs initiate testis development in response to disrupted oogenesis progression^2^. Therefore, we hypothesize that Rbpms2 negatively regulates several *rbtRNAs* in gonocytes to promote oocyte specification, oogenesis, and female sex differentiation.

**Figure 1.**
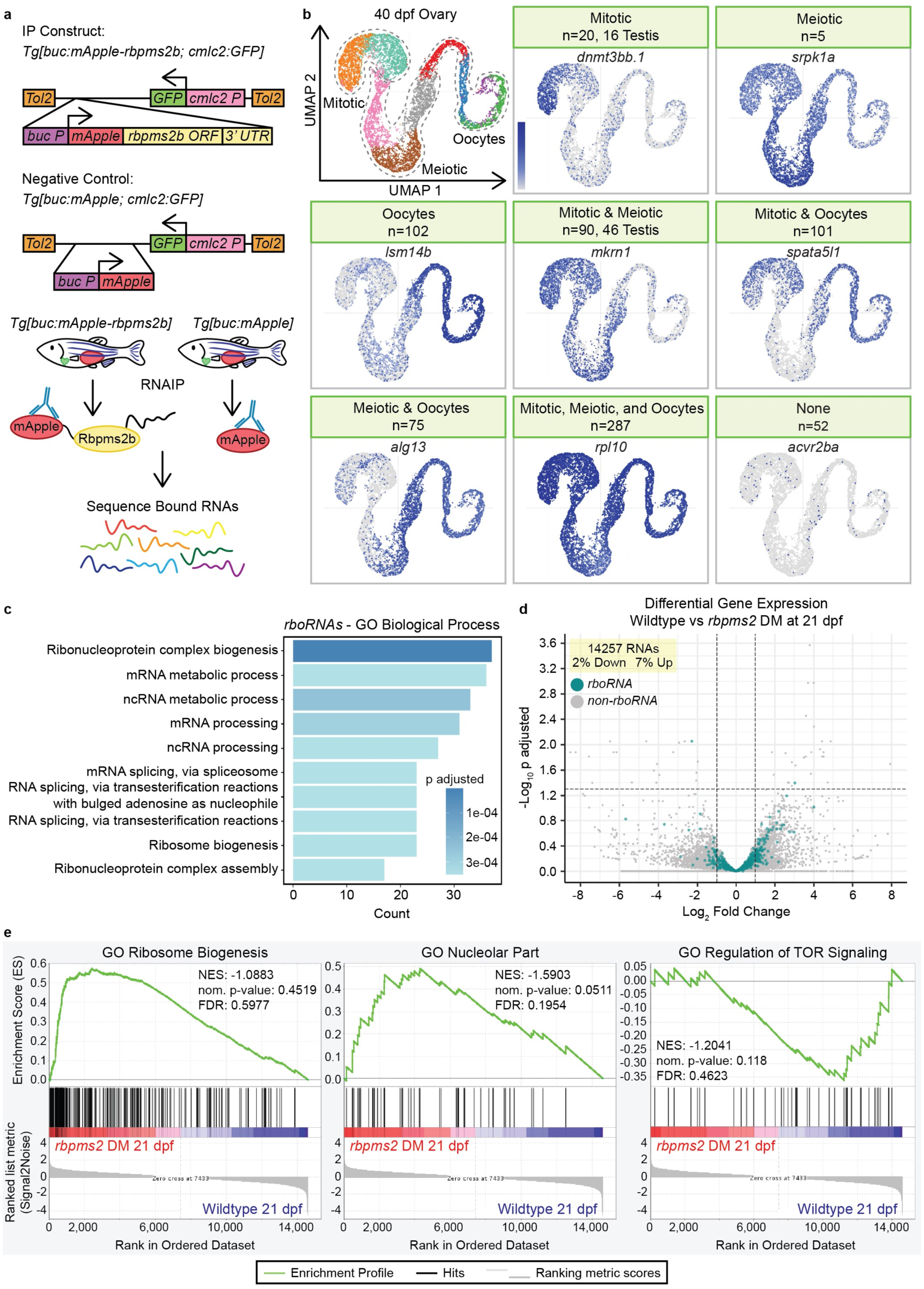
Rbpms2 binds and regulates RNAs required for testis development and ribosome and ribonucleoprotein biogenesis. (a) RNAIP and Negative control constructs and RNAIP scheme. (b) UMAP of cell populations in 40 dpf ovary^20^ and representative UMAPs of *rboRNAs* in a given cell population(s) of the 40 dpf ovary. Number of RNAs in a category are given in the header and *rboRNAs* associated with testis fates are specified in the header as well. (c) Top 10 GO biological processes associated with *rboRNAs*. (d) Volcano plot of differential gene expression in 21 dpf wildtype and *rbpms2* DMs from bulk RNA sequencing*. rboRNAs* are overlayed in blue. (e) Representative enrichment plots for GO Ribosome Biogenesis, GO Nucleolar Part, and GO Regulation of TOR Signaling from bulk RNA sequencing of 21 dpf wildtype versus *rbpms2* DMs.

Our previous work has shown that Rbpms2 is not only required to suppress testis fates but must also promote factors and pathways related to ovary fates for successful oogenesis^3^. To better understand the pathways by which Rbpms2 may promote ovary fates specifically, we performed GO Term Biological Process analysis on *rboRNAs* (Figure 1c). Of the Top 10 terms, several *rboRNAs* were related to mRNA metabolism and processing and ribosome biogenesis and ribonucleoprotein complex biogenesis and assembly. It has been established in invertebrates and vertebrates that a sufficient ribosomal quantity, and therefore translational capacity, is vital for productive oogenesis and subsequent embryogenesis (^25,26^, reviewed in^4^). This suggests that Rbpms2 may positively regulate *rboRNAs* related to ribosome biogenesis and ribonucleoprotein biogenesis and assembly to ensure sufficient levels of each are present for oogenesis.

Several mechanisms contribute to cell fate decisions, such as regulation of RNA biogenesis and metabolism, including RNA stabilization and decay which are known functions of RNAbps (reviewed in^27^), and are important for oogenesis^28–30^. To determine if Rbpms2 regulates *rboRNA* stability, we performed bulk RNA sequencing on *rbpms2* DMs and wild-type fish at 21 dpf, prior to sex determination, and evaluated global RNA and *rboRNA* differential expression. Of 14257 global RNAs present, only 2% were significantly downregulated and 7% upregulated between wildtype and *rbpms2* DMs (Figure 1d). Overlaying *rboRNAs* on this dataset showed that 728/730 targets are not significantly changed (Figure 1d). This suggests that Rbpms2 does not regulate RNA stability but could instead function at the level of translational control. Therefore, we conclude that 1) Rbpms2 likely functions to repress translation of *rbtRNAs* expressed in early gonocytes in ovaries and 2) promote translation of oocyte factors, including ribosome and ribonucleoprotein complex biogenesis regulators to support growth, oocyte differentiation, and meiotic progression.

To further understand the pathways by which Rbpms2 promotes oogenesis and female sex differentiation, we analyzed GSEA enrichment plots from our bulk RNAseq dataset (Figure 1e). RNAs related to ribosome biogenesis and the nucleolus, the hub of ribosome biogenesis, were highly enriched in *rbpms2* DMs as compared to wild-type fish. This further suggests that Rbpms2 likely functions at the level of translation to promote ribosome biogenesis, a process essential to oogenesis, and that the absence of Rbpms2 may disrupt a feedback loop between ribosome biogenesis, RNA transcription, and translation. Further, expression of RNAs related to regulation of TOR signaling are greatly decreased in *rbpms2* DMs at 21 dpf as compared to wildtype (Figure 1e). mTOR signaling, specifically via the mTORC1 nutrient-sensing branch, is a known regulator of nucleolar formation and all stages of ribosome biogenesis (reviewed in^31^). These results suggest that early gonocytes exist in a bipotential state wherein Rbpms2 mediates a binary fate-switch by repressing testis factors and promoting nucleolar formation and oocyte development via mTor signaling to ensure sufficient ribosome biogenesis and capacity for oogenesis.

### Rbpms2 regulates nucleolar amplification

Prophase I is a protracted phase of meiosis that occurs in early oocytes, and is a period of significant transcription prior to diplotene arrest (reviewed in^32^). During meiotic progression, oocytes cease transcription and ramp up translation to support major oocyte growth, prepare maternal reserves, and support the translational requirements of the future embryo. This translational demand requires a large pool of translational machinery, most notably the ribosome. Ribosome biogenesis occurs in the nucleolus, a structure composed of proteins, ribosomal DNA (rDNA), and ribosomal RNA (rRNA) assembled around nucleolar organizer regions into distinct compartments (reviewed in^33^). To support the high generation of ribosomes, teleosts and other fish and amphibian oocytes amplify their single nucleolus to multiple nucleoli in pachytene with nucleolar expansion and maturation culminating at diplotene when the oocyte arrests in meiosis I (Figure 2a)^34–36^.

**Figure 2.**
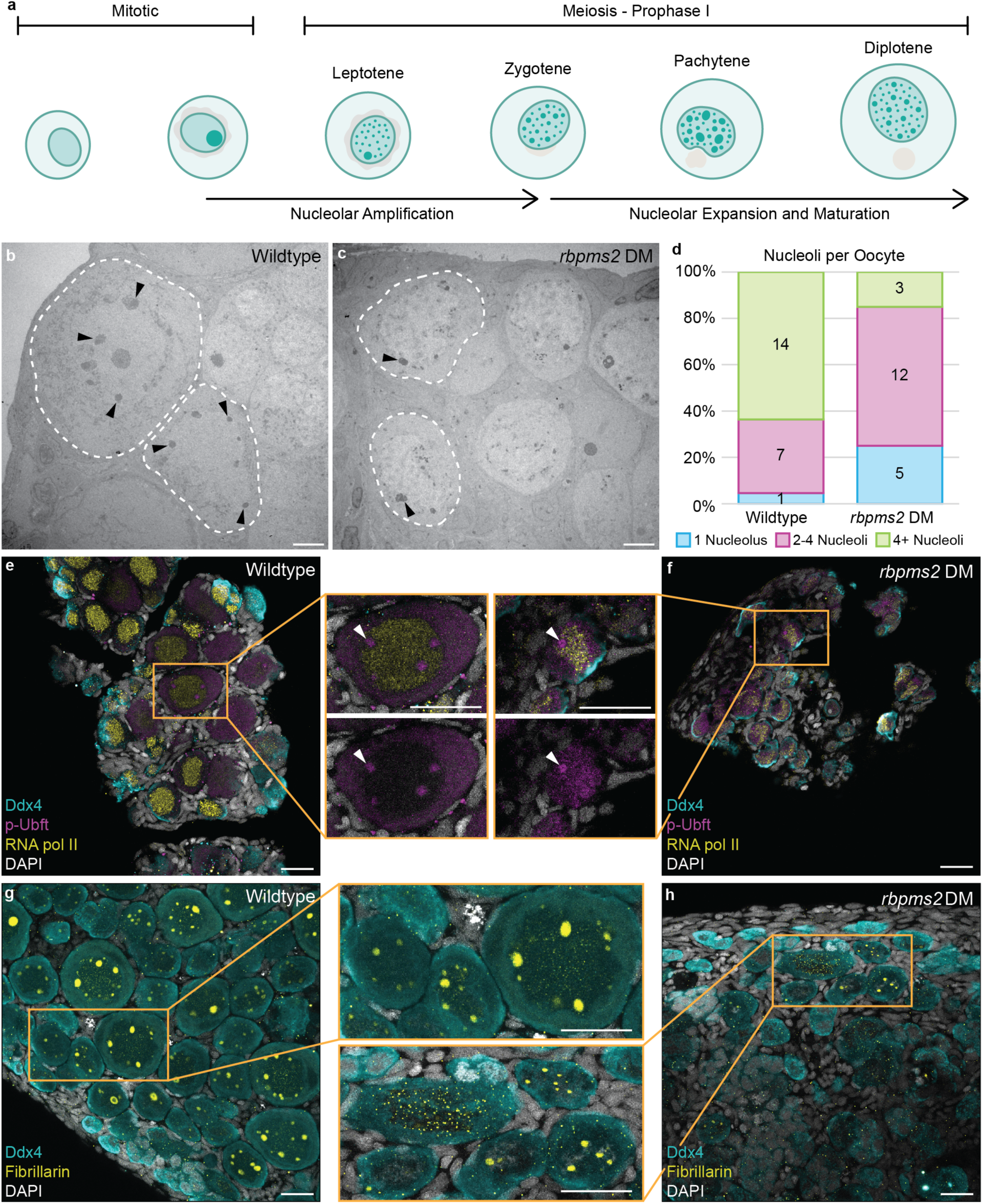
Nucleoli and ribosome biogenesis factors are dysregulated in *rbpms2* DMs. (a) Schematic of nucleolar development from mitosis to prophase I arrest. Briefly, upon entry into meiosis, the singular nucleolus of the germ cell disassembles. As the cell progresses through prophase I, new nucleoli are amplified through seeding by rDNA loci and nucleoli begin to expand. Nucleolar expansion, maturation, and ribosome biogenesis continues up until meiotic arrest at diplotene. (b-c) Transmission electron micrographs of 35 dpf wild-type and *rbpms2* DM oocytes. Dashed lines outline oocyte cytoplasmic membrane and black arrowheads indicate nucleoli. Scale bar is 5 µM. (d) Quantification of number of nucleoli per oocyte in wild-type (n=2 fish) and *rbpms2* DM (n=2 fish) gonads. A minimum of 20 cells per genotype were analyzed and categorized into the following groups: 1 nucleolus (blue), 2-4 nucleoli (purple), and 4+ nucleoli (green). (e-f) p-Ubf1 (active RNA pol I; purple) and RNA pol II (yellow) immunostaining in 31-35 dpf wild-type (*rbpms2a^ae30^; rbpms2b^sa9329^* HM; n=4) and *rbpms2* DM (n=4) germ cells. Enlarged views of boxed regions show representative localization of p-UBF1 in wild-type and *rbpms2* DM oocytes. Scale bar for all images is 20 µM. (g-h) Fibrillarin (yellow) immunostaining in 29-33 dpf wild-type (*rbpms2a^ae30^; rbpms2b^sa9329^* HM; n=4) and *rbpms2* DM (n=4) germ cells. Enlarged views of boxed regions show representative localization of Fibrillarin in wild-type and *rbpms2* DM oocytes. Scale bar for all images is 20 µM. (e-h) Germ cells are labelled by Ddx4 (teal) and nuclei are labelled with DAPI (white).

Given the abundance of nucleoli and ribosome biogenesis-related *rboRNAs*, we reanalyzed transmission electron microscopy (TEM) images of *rbpms2* DM oocytes to assess nucleolar features. Ultrastructural analyses and quantification of nucleoli number revealed that wild-type prophase I oocytes had greater than four nucleoli per cell at 35 dpf (Figure 2b, d) whereas most *rbpms2* DM oocytes had fewer than 4 nucleoli (Figure 2c, d). Nucleoli are composed of distinct compartments that regulate rRNA transcription, modification, or early ribosomal protein assembly (reviewed in^33^). To determine if nucleolar development requires Rbpms2, we analyzed markers of distinct ribosome biogenesis stages in wildtype and *rbpms2* DMs. RNA polymerase I (RNA pol I) is required for rRNA transcription and functions within the nucleolus. To visualize RNA pol I activity, we stained for phosphorylated Upstream binding transcription factor (p-Ubft), which is required for active RNA pol I. In wild-type ovaries, RNA pol I nuclear localization was exclusive to nucleoli, as expected, of all germ cells (Figure 2e). We also observed localization of RNA pol I in the cytoplasm of oocytes and early meiotic cells. To compare rRNA and mRNA transcription in wild-type oocytes, we assessed the distribution of RNA polymerase II (RNA pol II) and RNA pol I. In wild-type gonads, we observed RNA pol II in all mitotic and early meiotic germ cell nuclei in addition to some somatic cell nuclei. As expected, RNA pol II was strongly localized to oocyte nuclei and did not overlap with RNA pol I, which is restricted to nucleoli within the nucleus (Figure 2e, Supp. Figure 3a). In *rbpms2* DMs, while RNA pol I localized to the cytoplasm and nucleoli of mitotic germ cells, it was observed throughout oocyte nuclei and overlapped with RNA pol II nuclear localization (Figure 2f).

To further characterize nucleoli, we analyzed Fibrillarin, a conserved nucleolar small nuclear ribonucleoprotein component that localizes to nucleoli^37^. In wild-type ovaries, Fibrillarin localized to the single nucleolus of mitotic germ cells and was present in the nucleoli of oocytes of all stages (Figure 2g). In *rbpms2* DM oocytes, Fibrillarin was localized to a few, small nucleoli or dispersed throughout the nucleus with no specific localization (Figure 2h). Notably, nucleolar localization of RNA pol I and Fibrillarin were consistent with ultrastructural data indicating few nucleoli per oocyte in *rbpms2* DM ovaries. Taken together these results indicate that nucleolar recruitment of ribosome biogenesis factors is dysregulated in *rbpms2* DM oocytes and that Rbpms2 contributes to ribosome biogenesis and may do so by regulating translation of these factors to promote nucleolar expansion in primary oocytes.

### Rbpms2 functions upstream of the mTorc1 regulator, Mios

An essential pathway regulating nucleolar formation and ribosome biogenesis is the nutrient-sensing mTorc1 pathway^38^. Due to impaired nucleolar expansion in *rbpms2* DMs (Figure 2b-d) and dysregulated recruitment of ribosome biogenesis factors to nucleoli (Figure 2e-h), we determined if this phenotype was mTorc1-related. In wild-type ovaries, localization of phosphorylated p70-S6K (p-Ps6k), a kinase directly phosphorylated by the active form of mTorc1 (reviewed in^11^), was asymmetrically distributed in mitotic nuclei, diffusely localized in early meiotic cell nuclei, and shifted to the cytoplasm of oocytes (Figure 3a). In *rbpms2* DMs, while p-Ps6k localization in mitotic cells and gonocytes was comparable to wildtype, p-Ps6k was not detected in DM oocytes (Figure 3b). This observation suggests that dysregulated mTorc1 signaling in the absence of Rbpms2 contributes to impaired oogenesis.

**Figure 3.**
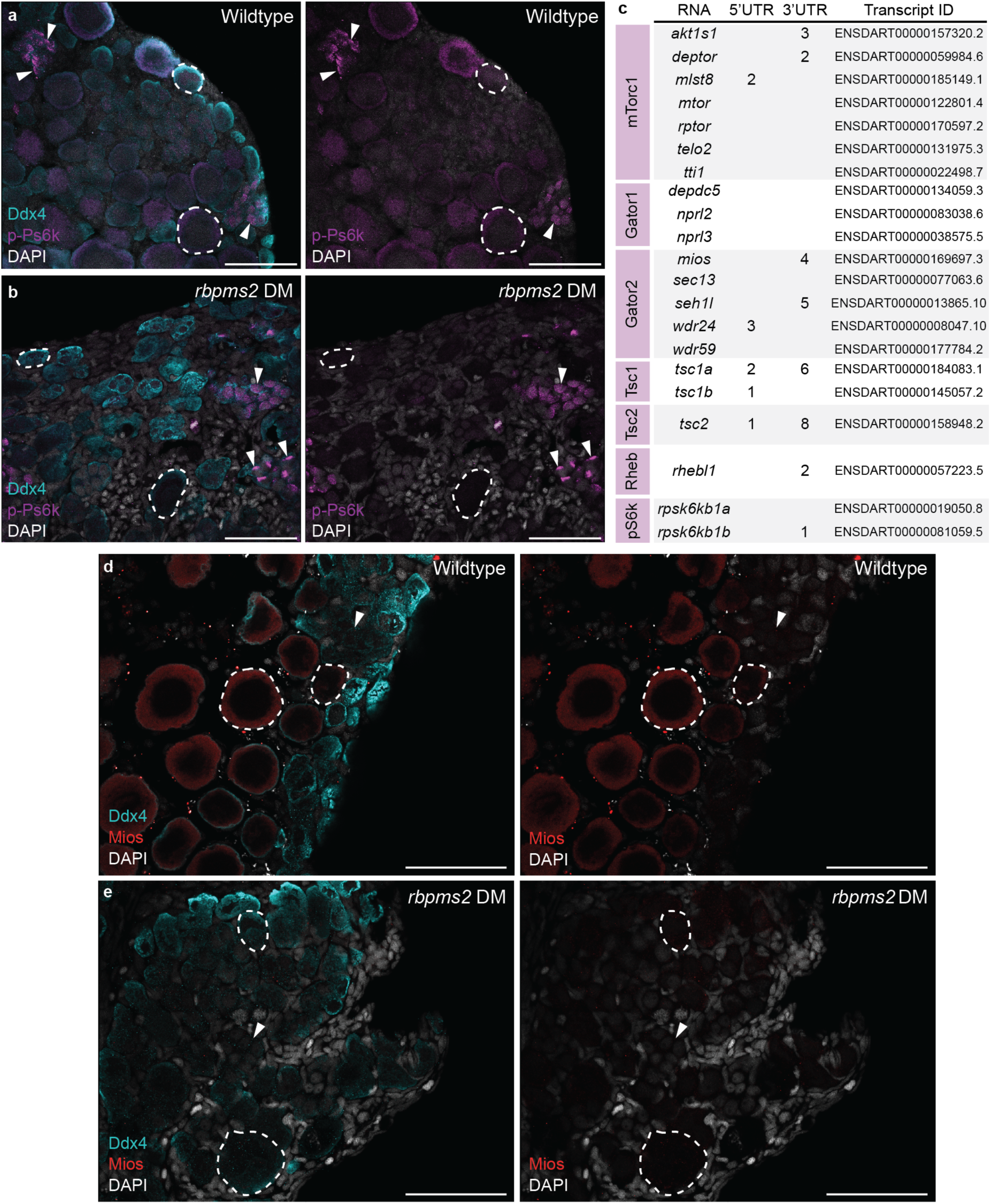
Rbpms2 functions upstream of the Gator2 complex protein, Mios. (a-b) Immunostaining of p-Ps6k (purple) in 29-33 dpf wild-type (*rbpms2a^ae30^; rbpms2b^sa9329^*HM; n=4) and *rbpms2* DM (n=4) germ cells. Germ cells are labelled with Ddx4 (teal) and nuclei are labelled with DAPI (white). Dashed lines indicate early oocytes and arrows indicate mitotic and early meiotic nuclei. Adjacent panel shows p-Ps6k and DAPI localization in specified cells. Scale bar is 50 µM. (c) Table of mTorc1-related components with Rbpms2 binding sites in their 3’ and/or 5’ UTRs. Transcript IDs evaluated for each component are listed and purple boxes indicate the protein component or complex of the pathway a given factor belongs to. (d-e) Immunostaining for Mios (red) in 29-31 dpf wild-type (*rbpms2a^ae30^; rbpms2b^sa9329^* HM, n=4) and *rbpms2* DM (n=4) germ cells. Germ cells are labelled with Ddx4 (teal) and nuclei are labelled with DAPI (white). Dashed outline indicates oocytes and adjacent panel shows Mios and DAPI localization in specified cells. Scale bar is 50 µM.

To investigate the relationship between Rbpms2 and mTorc1, we evaluated potential Rbpms2 targets related to amino acid-sensing and energy deficiency regulators of the mTorc1 signaling pathway (Figure 3c). Notably, the Gator2 complex protein, Missing oocyte (Mios), contains 4 Rbpms2 binding sites in its 3’ UTR and has been previously shown to be dispensable for viability but required for oogenesis in *Drosophila* ^18,39^. *mios* transcripts were detected in mitotic and meiotic germ cells of the 40 dpf ovary and was not differentially expressed between wildtype and *rbpms2* DMs at 21 dpf (Supp. Fig. 2a-b). However, Mios protein was only detected in wild-type oocytes (Fig 3d) but was not detected in *rbpms2* DM oocytes (Figure 3e). Together, these results suggest that Rbpms2 is a positive regulator of *mios* RNA translation and therefore acts upstream of mTorc1 signaling.

### Mios is required for nucleolar development and sustained oogenesis

To investigate the contribution of Mios to sex-specific germ cell differentiation in zebrafish, we obtained an ENU induced T>A point mutation from the Sanger Institute Mutation project, *mios^sa22946^,* which causes a premature stop codon (93L-> stop) within exon 2 (Figure 4a)^40^. We confirmed the *mios^sa22946^* mutation by sequencing both genomic DNA and cDNA and developed restriction digest based genotyping assays to distinguish the wild-type and mutant alleles (Figure 4a). In addition, we used CRISPR/Cas9 mutagenesis to generate additional mutant alleles disrupting exon 2 of zebrafish *mios*^41^. Several in-frame deletions were recovered including *mios^ms20^,* an 11bp deletion allele that leads to a frameshift and premature stop codon upstream of all Mios functional domains (Figure 4a). We confirmed that Mios protein is absent in *mios^sa22946^* mutant gonads; therefore, *mios^sa22946^* is likely a null allele (Supp. Figure 2c).

**Figure 4.**
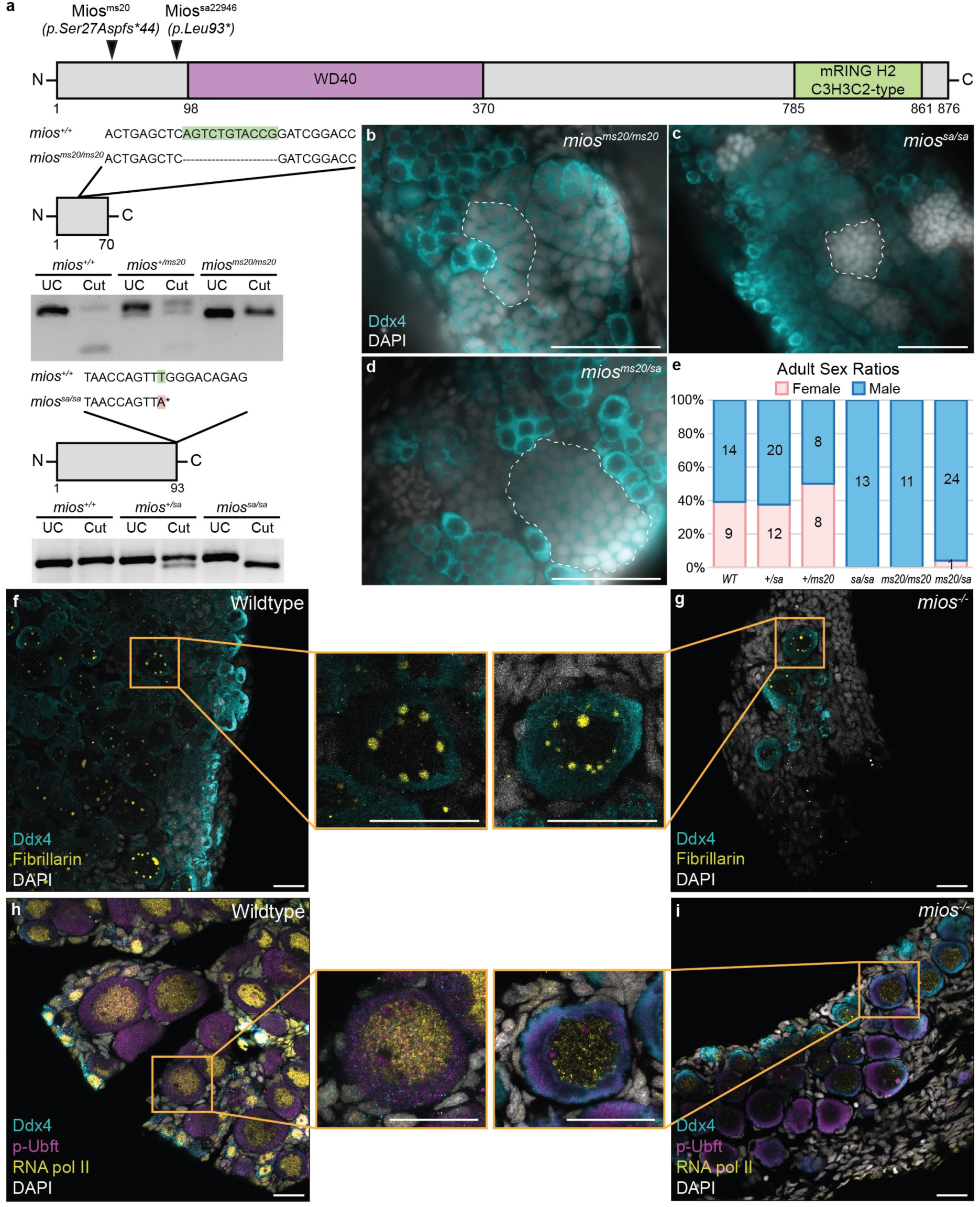
Mios is required for oogenesis and nucleologenesis. (a) Schematic of the Mios protein and mutagenized regions (black arrowheads) in the Mios^ms20^ and Mios^sa^ proteins. Truncated proteins are shown below with highlighted DNA mutations and corresponding genotyping assays. (b-d) Germ cell (Ddx4, teal) and DNA (DAPI, white) in *mios^ms20/ms20^* (50 dpf; n=5), *mios^sa/sa^* (45-51 dpf; n=4), and *mios^ms20/sa^*(50 dpf; n=4) fish. Developing sperm are outlined with dashed lines and scale bar is 50 µM. (e) Sex ratio graph for 60 dpf+ wildtype, heterozygous, and mutant fish. Pink represents female fish and blue indicates male fish. Number of fish screened are indicated for each group. (f-g) Fibrillarin (yellow) immunostaining in 35 dpf wild-type (n=7) and *mios^−/−^* (n=10) germ cells. Enlarged views of boxed regions show representative localization of Fibrillarin in wild-type and *mios^−/−^* oocytes. Scale bar for all images is 20 µM. (h-i) p-Ubf1 (active RNA pol I; purple) and RNA pol II (yellow) immunostaining in 35 dpf wild-type (n=4) and *mios^−/−^* (n=4) germ cells. Enlarged views of boxed regions show representative localization of p-Ubf1 (red) and RNA pol II (white) in wild-type and *mios^−/−^* oocytes. Scale bar for all images is 20 µM. (f-i) Germ cells are labelled by Ddx4 (teal) and nuclei are labelled with DAPI (white).

Analysis of *mios^ms20^* and *mios^sa22946^*heterozygous and mutant progeny showed they are viable to adulthood with no overt morphological deficits. However, homozygous mutants for *mios^sa22946^* grew slower than their heterozygous siblings, consistent with a deficit in mTorc1 signaling (Supp. Figure 2d). Additionally, gonad morphological and sex ratio analyses show that *mios^ms20^* and *mios^sa22946^* mutants and *mios^ms20/sa22946^* compound heterozygotes develop functional testes and differentiate as males, exclusively (Figure 4b-e). As *mios^ms20^*and *mios^sa22946^* gonad morphology and differentiation were indistinguishable, both alleles are likely protein null, and all further analyses were performed on *mios^sa22946^* fish (hereafter *mios^−/−^*). Investigation at earlier timepoints, specifically 35 dpf, showed that *mios^−/−^*initiate oogenesis. However, mutant oocytes remain small, and do not make it past diplotene, resulting in subsequent testis development (Figure 4c, g). Analysis of Fibrillarin in wild-type gonads revealed nucleolar amplification as expected (Figure 4f). In contrast, *mios^−/−^* oocytes contained a similar number of Fibrillarin but these puncta were much smaller (Figure 4g). Unlike Fibrillarin, RNA pol I showed no significant differences in nucleolar localization between wild-type and *mios^−/−^* oocytes (Figure 4h-i). In wildtype, exclusion of RNA pol II from nucleoli revealed a nucleolar region devoid of RNA pol I (Figure 4h). This is consistent with the localization of ribosome biogenesis factors to distinct compartments of nucleoli (reviewed in^33^). In *mios*^−/−^ oocytes, the nucleolar region devoid of RNA pol I was not observed (Figure 4i). This observation together with Fibrillarin staining indicates the nucleoli of *mios^−/−^* may be immature or have impaired recruitment of certain ribosome biogenesis factors.

To investigate nucleolar architecture, we performed ultrastructural analysis on wild-type and *mios^−/−^* oocytes (Supp. Figure 4a-b). We did not observe significant differences between nucleoli number, morphology, or size in wild-type and *mios^−/−^* oocytes (Supp. Figure 4a-d). This suggests that overt nucleolar architecture is not disrupted; however, nucleolar function, recruitment, or retention of specific factors may be impaired in the absence of Mios. Taken altogether, we conclude that impaired ovary differentiation is due to mutation of *mios* and that Mios, like Rbpms2, is required for oocyte differentiation in zebrafish.

### Mios promotes oogenesis through mTorc1

Next, we determined if Mios’ functions in oogenesis and female sex differentiation are mediated through mTorc1 signaling in ovaries. p-Ps6k localization in *mios^−/−^*was similar to early cells of wild-type gonads, however p-Ps6k in *mios^−/−^*oocytes was significantly reduced (Figure 5a-c; Supp. Figure 5a-b). To determine if activation of mTorc1 could suppress the *mios^−/−^* phenotype, we generated transgenic lines expressing constitutively active human L1460P mTOR (*mTOR^ca^*) under the germline-specific promoter, *ziwi*^42^ (Figure 5d). mTOR^ca^ has been shown to specifically increase mTORC1 downstream targets in HEK-293 cells^43^. We generated two alleles, *ms49* and *ms64,* of the *mTOR^ca^* transgene. Neither allele disrupted oogenesis or spermatogenesis in wild-type genotypes, and both restored female development in *mios^−/−^*, albeit to varying extents (Figure 5e). In fertility assays, *mios^−/−^*females expressing *mTOR^ca^* from transgenes had comparable fertility rates as their heterozygous and wild-type transgenic and non-transgenic female siblings (Figure 5f). However, *ms49* transgenic *mios^−/−^*females produced significantly more total eggs than wildtype non-transgenic siblings and significantly more degenerating eggs than all transgenic and non-transgenic siblings (Figure 5f, Supp. Figure 6a). While *ms64* transgenic *mios^−/−^*females did not differ from their transgenic and non-transgenic siblings in number of degenerating eggs produced, they produced a significantly higher number of total eggs as compared to their transgenic wildtype siblings (Figure 5f). These fish also produced several eggs with egg activation deficits (Supp. Figure 6b). These findings indicate incomplete suppression of oocyte loss or that Mios may have mTorc1 independent roles. These results demonstrate that the requirement for Mios in oogenesis is conserved. Further, in ovaries Mios acts through mTorc1 signaling and mTorc1 activation is sufficient for successful oogenesis in the absence of Mios.

**Figure 5.**
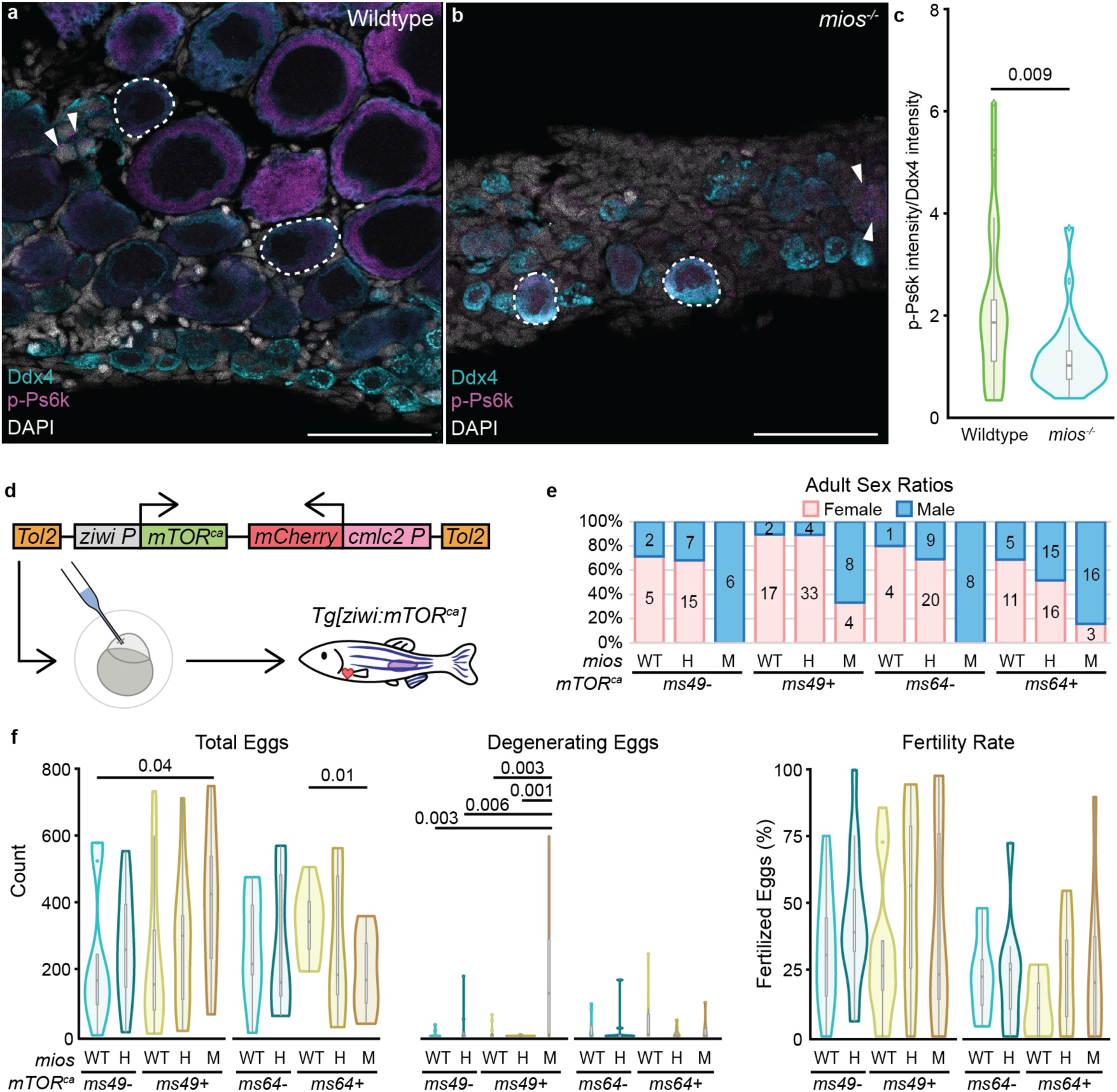
Mios promotes oogenesis through mTorc1 signaling. (a-b) Immunostaining of p-Ps6k (purple) in 35 dpf wild-type (n=7) and *mios^−/−^* (n=10) germ cells. Germ cells are labelled with Ddx4 (teal) and nuclei are labelled with DAPI (white). Dashed lines indicate early oocytes and arrows indicate mitotic and early meiotic nuclei. Adjacent panel shows p-Ps6k and DAPI localization in specified cells. Scale bar for (a,b) are 50 µM. (c) Quantification of the intensity of p-Ps6k normalized to the intensity of Ddx4 for wildtype (n=21 cells from 7 individual fish) and *mios^−/−^* (n=25 cells from 8 individual fish). Two-tailed paired equal variance student’s t tests were performed for the indicated groups with a significant p-value of less than 0.05. (d) *mTOR^ca^*transgene: the human *mTOR^ca^* expression is driven by the germline *ziwi* promoter and the heart specific promoter *cmlc2* drives *mCherry* expression as a selectable marker. Scheme depicts injection into 1-cell embryos and generation of stable *Tg[ziwi:mTOR^ca^]* lines. (e) Sex ratios for 90 dpf+ fish with *ms49* and *ms64 mTOR^ca^* alleles. Pink represents female and blue represents males; number of fish screened are indicated for each group. (f) Fertility assays and quantifications of total eggs, degenerating eggs, and fertility rates for *ms49* and *ms64 Tg[ziwi:mTOR^ca^]* fish. Two-tailed paired equal variance student’s t tests were performed for the indicated groups with a significant p-value of less than 0.05.

### Mios is required for oocyte differentiation independent of double strand break repair

In cancer cells, Gator2, inhibits Gator1, thus activating mTORC1^44^. In *Drosophila* ovaries, GATOR1 promotes meiotic entry of ovarian cysts by downregulating mTORC1 activity, which is subsequently reinforced by double strand breaks (DSBs)^18,45^. Accordingly, removal of GATOR1 or blocking DSB formation leads to elevated TORC1 activity, whereas removal of GATOR2 leads to persistent low mTORC1. Consistent with a model wherein GATOR2 opposes the activity of GATOR1 and DSBs, eliminating GATOR1 or DSBs suppresses *mio* mutant phenotypes in flies^18,39,45^. Spo11 (SPO11 initiator of meiotic double stranded breaks) is a conserved, essential enzyme for generating meiotic DSBs^46–48^. In zebrafish, Spo11 is not required for viability or oocyte production in females, although mutant gametes are likely aneuploid because embryos of mutant mothers are not viable^48^. To investigate if coordination between DSB repair and oocyte growth and progression through mTorc1 signaling is conserved, we generated double mutants lacking *mios* and *spo11* (*spo11^uc73^*)^48^. As expected, we saw no suppression or worsening of the *mios^−/−^*growth defect in the absence of *spo11* (Supp. Figure 2b). Although adult male and female *spo11^uc73/uc73^* were recovered, *mios^−/−^; spo11^uc73/uc73^* fish developed exclusively as males (Figure. 6a). Further, analyses of mutant gonads at 45 dpf revealed no delay or suppression of testis development in *mios*^−/−^; *spo11^uc7/uc73^* double mutants compared to *mios^−/−^* siblings (Figure. 6b-c). Based on these findings we conclude that eliminating DSBs is not sufficient to compensate for or bypass loss of Mios in zebrafish oocytes.

**Figure 6.**
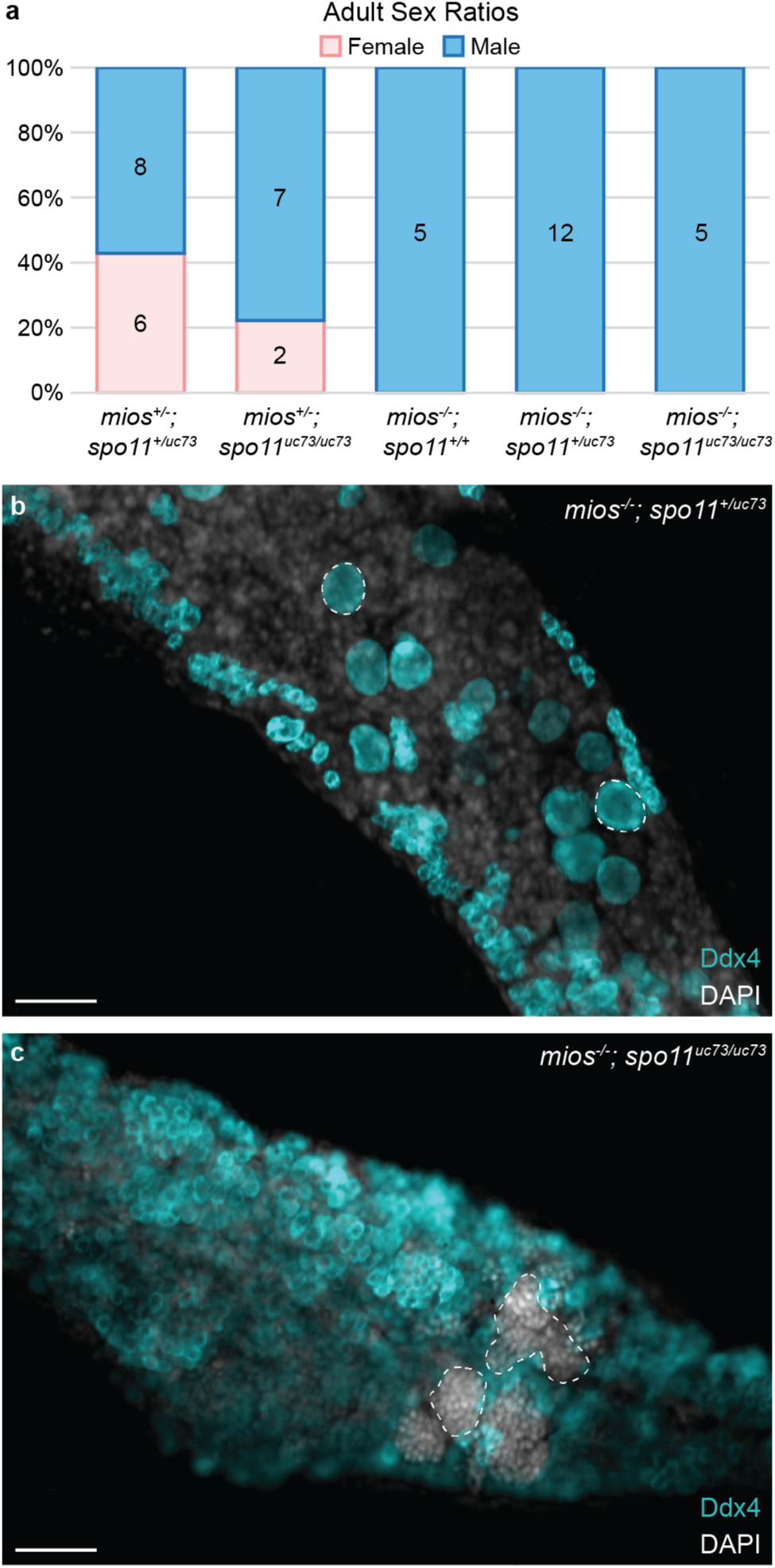
DSB inhibition does not restore oogenesis in *mios^−/−^*. (a) Sex ratio graph for 60 dpf+ *mios; tsc2^vu242^* HH, HW, and MH fish. Pink represents female fish and blue indicates male fish. Number of fish screened are indicated for each group. (b) Germ cell (Ddx4, teal) and DNA (DAPI, white) in *mios^−/−^* (*mios;spo11* MW/MH fish, n=6) and *mios^−/−^; spo11^uc73/uc73^*fish (n=7). Developing oocytes (b) and sperm (c) are outlined with dashed lines. Scale bar is 50 µM.

### mTorc1 activation in oogenesis requires Mios

If our hypothesis that failed differentiation of *mios^−/−^*oocytes is solely due to constitutive downregulation of mTorc1 activity, then activating mTorc1 signaling via modulation of the insulin/stress sensing arm of the pathway may suppress the *mios^−/−^* oocyte differentiation defect as occurs in *Drosophila*^18,45^. To investigate this possibility, we crossed the *mios^sa22946^* mutant allele onto the zebrafish *TSC complex subunit 2 (tsc2; tsc2^vu242^)* background^49^. Tsc2 functions in complex with TSC complex subunit 1 (Tsc1) to inhibit the activity of the mTorc1 activator, Rheb (reviewed in^10^). Although *tsc2^vu242/vu242^* fish are not viable to adulthood, *tsc2^+/vu242^* have been shown to have elevated mTorc1^49^. Thus, if oocyte arrest is due to diminished mTorc1 in *mios^−/−^*, then genetic reduction of *tsc2* might restore sufficient mTorc1 activity to allow for oocyte progression. However, comparison of *mios^−/−^; tsc2^+/+^* and *mios^−/−^; tsc2^+/vu242^* adult fish revealed no suppression of male development (Fig. 7a). To exclude the possibility that reduction of Tsc2 did not provide sufficient mTorc1 activation, we modulated Rheb to drive mTorc1 activation through the stress sensing arm. Specifically, we generated three independent stable transgenic lines expressing constitutively active rat S16H Rheb (*Rheb^ca^*)^50^ under the germline-specific *ziwi* promoter^42^ (Figure 7b). Germline expression of *Rheb^ca^* did not disrupt sex determination in wild-type genotypes nor prevent oocyte loss in *mios^−/−^* (Figure 7b). Since both Tsc2 reduction and Rheb^ca^ failed to suppress oocyte loss in *mios^−/−^*, we conclude that activating the metabolic stress arm of the mTorc1 pathway is not sufficient to support oocyte progression. Further, these results indicate progression beyond early oocyte development uniquely requires mTorc1 activation via the Gator2-Mios mediated nutrient sensing pathway arm.

**Figure 7.**
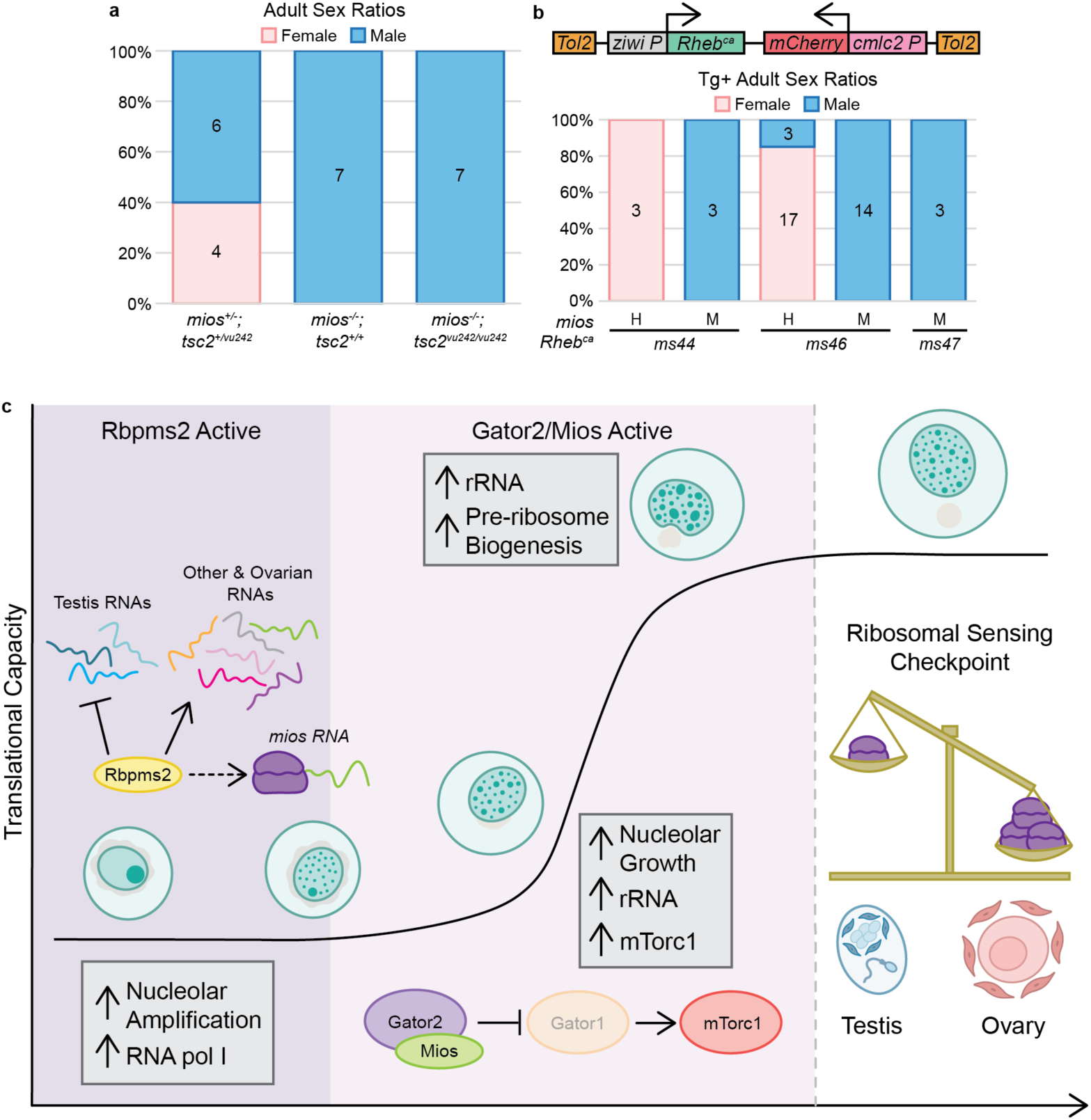
mTorc1 activation in oogenesis uniquely requires Mios. (a) Sex ratio graph for 60 dpf+ *mios; spo11^uc73^*HH, HM, MW, MH, and MM fish. Pink represents female fish and blue indicates male fish. Number of fish screened are indicated for each group. (b) *Rheb^ca^* transgene: the rat *Rheb^ca^* expression is driven by the germline *ziwi* promoter and the heart specific promoter *cmlc2* drives *mCherry* expression as a selectable marker. Sex ratios are presented for 60 dpf+ fish with the *ms44, ms46, and ms47 Rheb^ca^* alleles. Pink represents female and blue represents males; number of fish screened are indicated for each group. (c) In mitotic and early meiotic cells, Rbpms2 functions to repress translation of *rboRNAs* related to testis fates and promotes translation of *rboRNAs* related to mechanisms supporting oogenesis like ribosomal factors. Correspondingly, Rbpms2 functions upstream of nucleoli amplification and nucleolar localization of RNA pol I, which is required for rRNA transcription. In addition, Rbpms2 promotes translation of Mios. After sufficient nucleolar amplification, Gator2 and Mios activity increase mTorc1 signaling, and the nucleoli of differentiating oocytes expand and mature to support high levels of rRNA synthesis. As nucleoli grow, they develop an additional compartment that supports pre-ribosome biogenesis. Finally, as the cell progresses to diplotene, ribosomal abundance is measured. If sufficient ribosomes are present the cells continue through oogenesis. If a sufficient ribosome pool is not attained, then the gonocytes abort oogenesis and switch to spermatogenesis.

## DISCUSSION

Our study evaluates the mechanisms by which the vertebrate specific RNAbp, Rbpms2, promotes successful oogenesis and female sex determination and differentiation in zebrafish. Analyses of *rboRNAs* and global RNA changes indicate early oocytes express mRNAs encoding regulators of both sexual development programs and that Rbpms2 translationally represses testis-associated factors and promotes ovary-associated factors (Figure 7c). This is consistent with our previous findings that suppression of the male developmental program in *rbpms2* DMs prevents testis formation but is not sufficient to restore oogenesis^3^. We identify the mTorc1 pathway as an essential oogenesis pathway and a target of Rbpms2 activity, specifically through the Gator2 component, Mios. We find that Rbpms2 and Mios are both required for successful nucleolar development and consequently ribosome biogenesis, though they affect different stages of this process. Specifically, Rbpms2 acts upstream of nucleoli amplification and recruitment of the ribosomal RNA polymerase, RNA pol I, whereas Mios functions downstream. Additionally, we provide genetic evidence that the relationship between DSB regulation and mTorc1 signaling observed in *Drosophila* is not conserved in zebrafish oocytes. Finally, we demonstrate that the nutrient sensing arm of the mTorc1 pathway is uniquely required for oocyte progression and sustained oogenesis (Figure 7c). This finding is consistent with field observations linking nutrition availability to skewed sex ratios of developing zebrafish, specifically that nutrient deprivation causes male biased development^9^.

### Rbpms2 acts as a fate switch by binding and repressing translation of testis factors and promoting translation of ovarian factors

RNAbps are known to have broad, dynamic roles in post-transcriptional regulation in several cell types across species. Our previous work has demonstrated the role of Rbpms2 in regulating testis and ovary differentiation in zebrafish, but the mechanism and its targets remained to be determined^2,3^. Through RNAIP and bulk RNA-seq experiments, we provide evidence demonstrating the essential function of Rbpms2 as a multipronged regulator of sexual differentiation. We observe few changes in overall RNA and *rboRNA* transcript abundance between wildtype and *rbpms2* DMs. Additionally, RNA pol II, which we show is present in nuclei of wild-type zebrafish oocytes up to diplotene arrest, is also intact in *rbpms2* DM oocytes suggesting that Rbpms2 likely regulates sexual differentiation through translational control. Specifically, several *rboRNAs* are associated with testis functions and their expression is limited to the early, undifferentiated cell types of the 40 dpf ovary. This restricted expression is consistent with our hypothesis that Rbpms2 suppresses testis factors and promotes ovary factors as the gonad undergoes differentiation from a bipotential organ to an ovary. Moreover, several *rboRNAs* regulate a variety of processes, including those with defined roles in oogenesis and female sex differentiation. This further supports our previous finding that Rbpms2 must also positively regulate ovary-associated factors as silencing of testis differentiation pathways in *rbpms2* DMs did not restore or support oogenesis^3^.

Active antagonism of sex differentiation pathways has been observed both in organisms that undergo polygenic and sex chromosome-mediated sex determination mechanisms (reviewed in^51^). For example, in female mammalian sex determination, active suppression of SOX9 by FOXL2 is required for ovary formation and differentiation during embryonic development^52^. In many species the balance of androgens and estrogen levels is crucial to proper reproductive function. In humans, dysregulation of estrogen and androgen pathways contributes to a variety of female-associated pathologies, such as premature ovarian failure where androgen levels among other things are no longer maintained at appropriate levels for typical ovarian function^53^. Our findings position Rbpms2 as a crucial fate switch regulator of differentiation of the bipotential zebrafish gonad, supporting oogenesis and suppressing testis development by inhibiting testis-associated RNAs and promoting oogenesis related *rboRNAs*, affecting a wide array of downstream pathways.

### Rbpms2 promotes nucleoli development

Nucleoli serve as the hub for rDNA amplification, rRNA splicing, and pre-ribosome assembly (reviewed in^33^). In oocytes, extensive amplification of the nucleolus is required to support the growth demands of the oocyte as well as prepare the necessary components passed on from the oocyte to the embryo (reviewed in^54^). We show that several *rboRNAs* are related to ribosome biogenesis and the nucleolus and, more broadly, that these pathways are dysregulated between undifferentiated wild-type and *rbpms2* DM fish. Further, we find that nucleolar development is dysregulated in *rbpms2* DMs and *mios^−/−^* oocytes and likely contributes to failed ovary differentiation. The more severe nucleolar phenotypes and disruption of RNA pol I localization in *rbpms2* DMs but not *mios^−/−^* oocytes indicates differential contributions of the proteins to nucleologenesis and is consistent with Rbpms2 functions upstream of Mios. Specifically, our results suggest that rRNA transcription and nucleoli amplification require Rbpms2 or one of its targets, whereas Mios acts later in pre-ribosomal assembly (Figure 7c).

Transcription of rDNA has been posited to promote seeding of new nucleoli ^55^. Notably, in zebrafish it has been shown that demethylation and amplification of an rDNA locus at the end of chromosome four (*femrDNA*) strongly correlates with female sex determination and differentiation^56^. This suggests that oocyte development relies significantly on expansion of its ribosomal pool and positions ribosomal counting as a necessary checkpoint that must be passed for further oogenesis and female differentiation (Figure 7c). Our results indicate the importance of Rbpms2 in ribosome biogenesis. Further investigation is required to determine the relationship between Rbpms2 and *femrDNA* amplification in oocytes.

### Independent meiotic and gamete differentiation checkpoints in vertebrates

In *Drosophila*, Mio is required for oocyte progression beyond pachytene of prophase I and to maintain oocyte fate^39^. As *mio^−/−^* oocytes adopt the polyploid nuclear properties of nurse cells, oocyte fate is lost, resulting in female sterility^39^. Further, Mio loss in fly oocytes alters the kinetics of synaptonemal complex formation leading to delayed or abnormal synaptonemal complex formation^39^ such that oogenesis can be restored in *mio*^−/−^ by removing the *spo11* homolog, *mei-W86*^39,45^. As in *Drosophila,* zebrafish Mios is required in prophase I oocytes. However, unlike in flies, DSB sensing, and oocyte progression appear to be uncoupled in zebrafish, as loss of *spo11* in *mios^−/−^* gonads was not sufficient to restore oocyte differentiation and maturation. Further, *spo11^−/−^* zebrafish do not demonstrate a sex bias, have no overt growth defects, and progress through oogenesis and spermatogenesis (this study and ^48^). This is consistent with studies of mouse spermatogenesis, wherein inhibition of mTORC1 by conditional knockout of Raptor^57^ or rapamycin exposure^58^ did not disrupt DSB formation or repair. Moreover, *ex vivo* maturation of prematuration stage human and mouse oocytes in the presence of rapamycin results in reduction of yH2Ax, suggesting DSBs are either less numerous or are repaired faster in the context of mTOR inhibition^59^. Further evidence for distinct meiotic and nutritional oocyte differentiation checkpoints in vertebrates is failure of Tp53 loss to suppress oocyte loss in *rbpms2* DMs and other zebrafish mutants disrupting genes required for oocyte development through prophase I^2^ and of meiotic checkpoint factors, *e.g.* Mei-41/ATM/ATR, to suppress loss of Mio in *Drosophila*^39^. Collectively, the available evidence suggests that in vertebrates, the meiotic checkpoint and associated DNA integrity factors that safeguard faithful replication and reduction of chromosomes as part of meiosis operates independently of the mechanisms that ensure sex-specific gamete differentiation, which in oocytes is orchestrated in part by mTorc1 signaling.

## Methods

### Fish strains

Wild-type zebrafish embryos of the SAT strain were obtained from pairwise mating and were reared according to standard procedures^60^. Embryos were raised in 1X Embryo Medium at 28.5°C and staged accordingly^61^. *mios^ms20^* mutant fish were generated using Crispr-Cas9 mutagenesis as in as detailed below ^62^. The zebrafish *mios^sa22946^* allele was obtained from the Sanger Institute’s Zebrafish Mutation Project and acquired through Zebrafish International Resource Center (ZIRC)^40^. Complementation tests were performed by intercrossing carriers of *mios^sa22946^* and *mios^ms20^*. For epistasis analysis and to generate double mutants, *mios^sa22946^* heterozygous or mutant fish were crossed to fish with the *spo11^vc73^* or *tsc2^vu242^* allele, generated in ^48^ and ^49^, respectively. All procedures and experimental protocols were performed in accordance with NIH guidelines and were approved by the Icahn School of Medicine at Mount Sinai Institutional (ISMMS) Animal Care and Use Committees (IACUC #2017-0114).

### Mutagenesis

The zebrafish *mios^sa20^* allele was generated by CRISPR-Cas9 mediated mutagenesis^62^. *mios* single-guide RNAs (sgRNA) targeting exon 2 were designed using the CHOPCHOP webtool^63^. Briefly, the gene specific target and the constant oligonucleotides were annealed, and the overhangs were filled in using T4 DNA polymerase. In vitro transcription of the sgRNA was performed using the MEGAscript SP6 kit (Life technology, Ambion). 1 nL of 12.5 ng/µL of sgRNA and 1 nL of Cas9 protein (300 ng/µL)^64^ along with phenol red (Sigma Aldrich) were co-injected in one-cell stage embryos. At 24 hpf, 8 injected embryos and 8 control uninjected siblings were assayed by PCR amplification and T7 endonuclease (New England Biolabs, M0302S) digest for mutation analysis^41^. Individuals with new gel banding patterns were sequenced to characterize induced mutations. Injected embryos were raised to adulthood and screened for germline mutations by extracting genomic DNA from their progeny and fin tissue and assaying as above. The presence of digested fragments compared to the wild-type allele indicated *de novo* mutations. The corresponding bands were extracted from the gel, cloned into a PCR4 TOPO vector (Invitrogen, K457502), and sequenced in both directions to determine the induced mutation. Fish harboring *mios^ms20^* mutations were outcrossed to SAT fish or crossed to *mios^sa22946^* for complementation tests. All mutations were verified by sequencing both genomic DNA (gDNA) and cDNA from mutant animals. Total RNA was extracted from individual gonads from heterozygote intercrosses or SAT wildtype, using TRIzol (Life Technologies, 15596). The SuperScript III/IV Reverse Transcription Kit (Life Technologies, 18080-051) was used to prepare cDNA, and Easy-A High Fidelity Taq polymerase (Agilent, 600400) was used for RT-PCR to amplify the *mios* coding region. Wildtype and mutant PCR fragments were TOPO cloned into pCR8/GW/TOPO (Invitrogen, K250020) and sequenced (Psomagen). Sequences were analyzed using SnapGene software. All oligos used are in Supplemental Table 1.

### Genotyping

gDNA was obtained either from embryos, dissected juvenile and adult trunks, or fin tissue. Samples were lysed in an alkaline lysis buffer (25 mM NaOH, 0.2 mM EDTA, pH12) and heated at 95°C for 20 minutes. After cooling to 4°C, neutralization buffer was added (20 mM Tris-HCl, 0.1 mM EDTA, pH 8.1)^65^. The genomic region around *rbpms2a^ae30^* was amplified for 35 cycles with an annealing temperature of 60°C and the wild-type allele was digested with HaeIII for 1 hour (New England Biolabs, R0108S)^2^. The *rbpms2b^sa9329^* surrounding genomic region was amplified for 35 cycles with an annealing temperature of 60°C and the mutant allele was digested with MboII (New England Biolabs, R0148S)^2^ or the wild-type allele was digested with HphI (New England Biolabs, R0158S) for 1 hour. The region around the *mios^sa22946^* allele was amplified for 35 cycles using 60°C for annealing and the mutant allele was identified by restriction enzyme digestion with DdeI for 1 hour (New England BioLabs, R0175S). The genomic region around the *mios^sa20^* allele was amplified for 35 cycles using 57°C for annealing and identified by restriction enzyme digestion of the wild-type allele with BsaWI for 1 hour (New England BioLabs). The genomic region surrounding the *spo11^uc73^*allele was amplified for 35 cycles with an annealing temperature of 57°C and products were resolved without digestion^48^. The surrounding *tsc2^vu242^* genomic region was amplified for 35 cycles with an annealing temperature of 57°C and the wild-type allele was digested with HpyCH4IV for 1 hour (New England Biolabs, R0169S). Undigested and digested products were resolved in a 3% Metaphor 1:1 (Lonza)/agarose gel (Invitrogen) gel. All genotyping primers are listed in Supplemental Table 1.

### Dissections and length measurements

Fish were anesthetized with a lethal dose of tricaine (MS-22). Fish were positioned laterally, and rostral-caudal body length was measured prior to dissection. Trunks and gonads were prepared by removal of the head and tail regions, exposure of the body cavity by opening the ventral body wall, and removal of the gut tissue while leaving the gonads intact. Alternatively, gonads were dissected from the trunk and reserved. Trunks and gonads were preserved by flash freezing in liquid nitrogen and storage at −80°C or fixation as described below.

### Western blot

Individual gonads were dissected, lysed in 40 uL RIPA lysis buffer with protease inhibitor (Thermo Scientific, A32955) for 30 minutes at 4°C, vortexed, and centrifuged at 4°C. Lysates were removed and stored overnight at −80°C. Sample lysates (18 uL) were then combined with equal volumes 2x SDS loading buffer and boiled for 5 minutes prior to loading. 20 μl per sample were loaded on a 4-12% SDS-PAGE gel and proteins were transferred to PVDF membranes. Membranes were blocked in 5% milk in 1x PBS for 1 hour at room temperature. Anti-Mios antibody (Cell Signaling, 13557) was used at 1:250 and incubated overnight at 4°C. Membranes were washed for 5 minutes 3 times in 1x TBS-Tween and then for 5 minutes twice in TBS. Rabbit-HRP secondary antibody (Cell Signaling, 7074) was diluted 1:5000. Following a 1 hour incubation at room temperature, membranes were washed for 5 minutes 3 times 1x TBS-Tween and then for 5 minutes twice in 1x TBS. Proteins were detected with an ECL-Plus kit (Cytiva Lifescience, RPN2232). Chemiluminescence was imaged using a BioRad imager.

### Immunohistochemistry

For whole-mount immunofluorescence staining of gonads, tissues were fixed in 4% paraformaldehyde (PFA) overnight at 4°C. The next day, the samples were washed in 1x PBS,dehydrated in 100% methanol (MeOH) overnight, and stored at −20°C until use. Samples were gradually rehydrated with 1x PBS prior to acetone permeabilization at −20°C for (1) 7 minutes and blocked in 1x PBSTw (0.1% Tween20 in 1x PBS) with 5% Normal Goat Serum and 2% dimethyl sulfoxide (DMSO) for 4 hours minimum at 25°C or (2) for 2 minutes and blocked in 1x PBSTw with 2% Normal Goat Serum and 2mg/mL Bovine Serum Albumin for 4 hours minimum at 25°C. To stain for germ cells, samples were blocked (1 or 2) and the chicken anti-Vasa (Ddx4) antibody^48^ was used at a 1:3000 dilution and incubated at 4°C for 24 to 48 hours. Nucleolar labeling and mTorc1 activity assessments were accomplished by blocking samples (1) and incubating with mouse anti-Fibrillarin (1:750 dilution; Novus Biologicals, NB300-269), rabbit anti-p-p70s6k (1:500 dilution; Cell Signaling, 9205), rabbit anti-Mios (1:500 dilution; Cell Signaling, 13557) at 4°C for 24-48 hours. RNA pol I and II visualization was accomplished by blocking (2) and incubating samples with rabbit anti-p-UBF1 (1:100 dilution; Invitrogen, PA5-105512) and mouse anti-RNA pol II (1:100 dilution; Santa Cruz, sc-56767) and incubated at 4°C for 30 hours minimum.

For all experiments, samples were washed with 1xPBSTw several times and incubated at 4°C for 24-48 hours with chicken Alexa Fluor 488, rabbit Alexa Fluor 568, and mouse Alexa Fluor 647 (Molecular Probes) secondary antibodies diluted 1:500 in (1) or 1x PBSTw (samples initially blocked in (2)). Samples were then washed in 1x PBSTw several times, dissected, and mounted in Vectashield with 4′,6-diamidino-2-phenylindole (DAPI) (Vector Laboratories), Prolong Diamond Antifade Mountant with DAPI (Invitrogen), or incubated briefly (30 seconds to 15 minutes minutes) in DAPI, washed with 1x PBSTw, and mounted in Prolong Diamond Antifade Mountant (Invitrogen). Images were acquired using a Zeiss Zoom dissecting scope equipped with Apotome.2, Zeiss Axio Observer inverted microscope equipped with Apotome.2 and a charged-coupled device camera, or a confocal microscope Zeiss 880 AiryScan2.0 (40x or 63x oil objective, 1 µM stack increments, 1024 × 1024 pixel format) at the Microscopy CoRE at ISMMS. Image processing was performed in Zen Blue (Zeiss), ImageJ/FIJI, and Adobe Illustrator.

### Fluorescence Quantification

Fluorescence was quantified using ImageJ/Fiji by determining the area and mean intensity of cytoplasmic Ddx4 and the identical region occupied by p-Ps6k. The average intensity for each was determined by dividing the mean intensity by the area and the average fluorescence intensity of p-Ps6k was then normalized to the average fluorescence intensity of Ddx4. Fluorescence was quantified from average intensity projections of 3, 1 µm slices and a minimum of 1 oocyte was quantified per fish.

### RNA-IP

Wildtype ovaries were dissected from 90 dpf+ adult females from stable lines expressing oocyte-specific mApple tagged Rbpms2 protein (*Tg[buc:mApple-Rbpms2b-3’UTR]*; n=2), or an oocyte-specific mApple only (*Tg[buc:mApple]*; n=2) control line^2^, snap frozen, and stored at −80°C. We performed RNAIP adapted from previously published methods^23,66^. Briefly, ovaries were crosslinked in 250 µL YSS buffer (50 mM pH 8.0 Tris HCl, 50 mM NaCl, 0.1% NP40, 1 U/mL of RNAsin (Roche, 3335402001), protease inhibitor (Thermo Scientific, A32955), 0.5 mM DTT, 100 mM Sucrose) for 10 seconds in a UV crosslinker at 400mJx100cm^2^ then homogenized. 750 µL YSS buffer was added to each homogenate and centrifuged. Pellets were resuspended in 1 mL YSS buffer. 250 µL of resuspended pellet was pre-cleared with 30 µL pre-washed protein G Dynabeads (Invitrogen, 10004D) for 1 hour at 4°C. 25 µL primary anti-RFP antibody (ChromoTek, 5f8) diluted 1:500 in YSS was added to each 250 µL pre-cleared lysate and incubated overnight at 4°C with rocking. After incubation, 30 µL pre-washed Dynabeads were added to each sample and rocked at 4°C for 2 hours. At 4°C, magnetic beads were washed five times with YSS buffer, resuspended in 100 µL Proteinase K lysis buffer (10 mM Tris HCl pH 8.0, 50 mM NaCl, 5 mM EDTA, 0.5% SDS) with 10 µg proteinase K (New England Biolabs, P8107S), vortexed briefly, and incubated at 50°C for 1 hour. Dynabeads were then removed, RNA was isolated using TRIzol (Life Technologies, 15596), and precipitated with 3M sodium acetate. Precipitated RNA was stored at −80°C until RNA library prep.

### RNAseq

For sequencing of RNAIP isolated RNAs, samples were processed (Qiagen, 74104) and cytoplasmic and mitochondrial rRNA-deleted stranded Total RNA libraries were prepared using Illumina Truseq RNA Library Prep kit (Illumina 20020589) from four individual adult ovaries (2 *buc:mApple-Rbpms2b-3’UTR* crosslinked and 2 *buc:mApple* crosslinked). Libraries were sequenced using an Illumina MiSeq, ∼150bp read length, paired ends, with ∼ 1 million sequencing reads per sample. Sequencing data was aligned to the zebrafish genome GRCz10 using the Illumina website. RNAs pulled down with mApple tagged Rbpms2 and mApple alone were compared from the cross-linked and uncrosslinked samples. Only RNAs that appeared in both the cross-linked and uncrosslinked mApple-Rbpms2, but not in controls were considered Rbpms2 target RNAs.

For bulk RNA-sequencing, individual gonads were dissected at 21 dpf from *rbpms2* DMs (n=4) and siblings (*rbpms2a^ae30^; rbpms2b^sa9329^* WM n=4) and flash frozen in 100 µL of fresh RNA lysis buffer^23^ containing 2-mercaptoethanol and stored at −80°C prior to RNA purification. RNA was extracted and purified from individual tissues according to kit instructions (Agilent Absolutely RNA Nanoprep Kit, 400753), eluted in 13 µL RNA elution buffer, and stored at −20°C prior to sequencing. RNA quality and quantity was assessed on an Agilent 2100 Bioanalyzer chip (RIN>8). Libraries were generated and sequencing was performed by the NY Genome Technology Center with a low input SMART-Seq HT with Nxt HT kit (Clontech Laboratories, 634947) and SP100 cycle flow cell. All samples were sequenced on an Illumina SovaSeq 6000, paired end reads. Analysis of Fastq files and differential gene expression analyses were performed using BasePair pipelines. Further analyses were performed in R (v4.2.2). GO Term Biological Process Analysis was performed using clusterProfiler^67,68^, GOfuncR^69^, GOplot^70^, genekitr^71^, and rrvgo^72^ packages and the volcano plot was generated using the EnhancedVolcano^73^ package. Corresponding plots were generated with the ggplot2^74^ package.

### Transgenesis

Generation of the *Tg[Tol2-ziwi:mTOR^ca^; cmlc2:mCherry-Tol2]* and *Tg[Tol2-ziwi:Rheb^ca^; cmlc2:mCherry-Tol2]* constructs was accomplished by using Gateway Recombination multiple-fragment cloning with the LR II+ Cloning Enzyme Mix (Invitrogen, 12538120). The *mTOR^ca^* and *Rheb^ca^* sequences (pME-mTOR L1460P^43^; pME-Rheb S16H^50^) were recombined with p5E-ziwiP^42^ into the pBH-R4/R2 destination vector^23^. 1-cell embryos were injected with 1 nL of plasmid solution (100 ng of plasmid, 2.5 µL of Tol2 transposase RNA transcribed from pCS2FA-transposase, 0.25 µL of phenol red (Sigma Aldrich), and water (up to 5 µL)) and screened for mCherry heart expression at 1-3 dpf. Progeny of these fish were then screened for germline transmission of the given transgene by mCherry heart expression and stable lines were propagated.

### Fertility Assay

Female transgenic (*mios* wildtype n=3, heterozygous n=3, and mutant n=3) siblings and non-transgenic (*mios* wildtype n=3 and heterozygous n=3) siblings for both the *ms49* and *ms64 mTOR^ca^*alleles were crossed to transgenic male siblings, unrelated non-transgenic *mios* heterozygotes or mutants, or non-transgenic unrelated SAT fish weekly for 4-5 (*ms49* fish) or 3 (*ms64* fish) crosses total. All eggs were collected within two hours of initial mating and the number of total (degenerating + fertilized + unfertilized), degenerating, fertilized, and unfertilized eggs were counted. Fertility rate was calculated by dividing the number of fertilized eggs by the total number of eggs for a given cross and multiplying by 100.

### Electron Microscopy

2 wild-type and 2 *rbpms2* DM 35 dpf fish were euthanized in tricaine (MS-22) and gonads were dissected and processed as previously described^2^. 4 *mios^+/-^*and 4 *mios^−/−^* 35 dpf fish were euthanized in tricaine (MS-22),dissected, and immediately fixed in Karnovsky’s solution (2% glutaraldehyde and 2% paraformaldehyde in 0.1 M sodium cacodylate) for at least 24 hours at 4°C. The samples were then washed with cacodylate buffer, osmicated with 1% osmium tetroxide for 1 hour, and en bloc stained with 2% uranyl acetate for 1 hour. After a quick rinse in water and seven 10-min incremental ethanol dehydration steps (25%-100%), samples were infiltrated with a mixture of propylene oxide and an epoxy resin (Epon, Electron Microscopy Sciences). Samples were polymerized in pure resin in a vacuum oven at 60°C for 72 hours. Ultrathin sections were cut at 90 nm using a diamond knife (Diatome) on an ultramicrotome (Leica EM UC7) until the majority of cell nuclei were visible and mounted onto a formvar-supported slot grid (Electron Microscopy Sciences). Sections were imaged using a Hitachi H7500 TEM at 75 kV and 2048×2048 pixel, and 16-bit images of at least 10 oocytes per sample were taken using a CCD camera (AMT Imaging). Images were analyzed with FIJI/ImageJ and the Biodock Alligator model architecture^75^.

## Supporting information

Wilson et al Supplemental Materials

## Acknowledgements

We thank the members of the Marlow and Rangan labs for helpful discussions and The Center for Comparative Medicine staff at ISMMS for fish care. Microscopy was performed at the Microscopy and Advanced Bioimaging CoRE at the Icahn School of Medicine at Mount Sinai. Bulk RNA sequencing was performed at the New York University Langone’s Genome Technology Center (RRID: SCR_017929). We thank the Zebrafish International Resource Center for providing fish lines and Dr. Wenbiao Chen for providing the *mTOR* L1460P and *Rheb* S16H pME constructs. This work is supported by startup funds to F.L.M. and the National Institutes of Health (R01-GM089979), M.L.W. (1F31HD112158-01), S.R. (1F32HD097898-01A1), and O.K. (T32-GM007288, F30HD082903). S.R. was supported by a New York Stem Cell Foundation training grant (C32561GG). Work in the Draper lab was supported by R01 HD-081551 and NSF/IOS-1456737 to B.W.D.

## Author Contributions

Conceptualization: M.L.W, S.R., O.K., and F.L.M. Methodology: M.L.W., S.R., Y.L., O.K., B.W.D., and F.L.M. Investigation: M.L.W, S.R., N.K., D.A., Y.L, O.K., F.L.M. Visualization: M.L.W, S.R., N.K., D.A., Y.L, O.K, B.W.D., and F.L.M. Supervision: B.W.D. and F.L.M. Writing—original draft: M.L.W. and F.L.M. Writing—review and editing: M.L.W, S.R., N.K., D.A., Y.L, O.K, B.W.D., and F.L.M.

## Competing Interests

The authors declare that they have no competing interests.

## Notes

### Competing Interest Statement

The authors have declared no competing interest.

## References

1 Von Stetina, J. R. & Orr-Weaver, T. L. Developmental control of oocyte maturation and egg activation in metazoan models. Cold Spring Harb Perspect Biol 3, a005553 (2011). 10.1101/cshperspect.a005553

2 Kaufman, O. H., Lee, K., Martin, M., Rothhamel, S. & Marlow, F. L. rbpms2 functions in Balbiani body architecture and ovary fate. PLoS Genet 14, e1007489 (2018). 10.1371/journal.pgen.1007489

3 Romano, S., Kaufman, O. H. & Marlow, F. L. Loss of dmrt1 restores zebrafish female fates in the absence of cyp19a1a but not rbpms2a/b. Development 147 (2020). 10.1242/dev.190942

4 Mercer M, J. S., Ni C, Buszczak M. The Dynamic Regulation of mRNA Translation and Ribosome Biogenesis During Germ Cell Development and Reproductive Aging. Front Cell Dev Biol. 9, 1–23 (2021). 10.3389/fcell.2021.710186

5 Gu, L. et al. Metabolic control of oocyte development: linking maternal nutrition and reproductive outcomes. Cell Mol Life Sci 72, 251–271 (2015). 10.1007/s00018-014-1739-4

6 Lopez, A. L., 3rd et al. DAF-2 and ERK couple nutrient availability to meiotic progression during Caenorhabditis elegans oogenesis. Dev Cell 27, 227–240 (2013). 10.1016/j.devcel.2013.09.008

7 Igosheva, N. et al. Maternal diet-induced obesity alters mitochondrial activity and redox status in mouse oocytes and zygotes. PloS one 5, e10074 (2010). 10.1371/journal.pone.0010074

8 Drummond-Barbosa, D. & Spradling, A. C. Stem cells and their progeny respond to nutritional changes during Drosophila oogenesis. Developmental biology 231, 265–278 (2001). 10.1006/dbio.2000.0135

9 Lawrence, C., Ebersole, J. P. & Kesseli, R. V. Rapid growth and out-crossing promote female development in zebrafish (Danio rerio). Environmental Biology of Fishes 81, 239–246 (2007). 10.1007/s10641-007-9195-8

10 Takahara, T., Amemiya, Y., Sugiyama, R., Maki, M. & Shibata, H. Amino acid-dependent control of mTORC1 signaling: a variety of regulatory modes. J Biomed Sci 27, 87 (2020). 10.1186/s12929-020-00679-2

11 Sengupta, S., Peterson, T. R. & Sabatini, D. M. Regulation of the mTOR complex 1 pathway by nutrients, growth factors, and stress. Mol Cell 40, 310–322 (2010). 10.1016/j.molcel.2010.09.026

12 Guo, J. et al. Oocyte stage-specific effects of MTOR determine granulosa cell fate and oocyte quality in mice. Proc Natl Acad Sci U S A 115, E5326–E5333 (2018). 10.1073/pnas.1800352115

13 Bandyopadhyay, A., Bandyopadhyay, J., Chung, J., Choi, H. S. & Kwon, H. B. Inhibition of S6 kinase by rapamycin blocks maturation of Rana dybowskii oocytes. Gen Comp Endocrinol 113, 230–239 (1999). 10.1006/gcen.1998.7199

14 Morley, S. J. & Pain, V. M. Hormone-induced meiotic maturation in Xenopus oocytes occurs independently of p70s6k activation and is associated with enhanced initiation factor (eIF)-4F phosphorylation and complex formation. J Cell Sci 108 (Pt 4), 1751–1760 (1995). 10.1242/jcs.108.4.1751

15 Fukuyama, M. et al. C. elegans AMPKs promote survival and arrest germline development during nutrient stress. Biol Open 1, 929–936 (2012). 10.1242/bio.2012836

16 Jeong, E. B., Jeong, S. S., Cho, E. & Kim, E. Y. Makorin 1 is required for Drosophila oogenesis by regulating insulin/Tor signaling. PloS one 14, e0215688 (2019). 10.1371/journal.pone.0215688

17 Gorre, N. et al. mTORC1 Signaling in oocytes is dispensable for the survival of primordial follicles and for female fertility. PloS one 9, e110491 (2014). 10.1371/journal.pone.0110491

18 Wei, Y. et al. TORC1 regulators Iml1/GATOR1 and GATOR2 control meiotic entry and oocyte development in Drosophila. Proc Natl Acad Sci U S A 111, E5670–5677 (2014). 10.1073/pnas.1419156112

19 Mayer, C. & Grummt, I. Ribosome biogenesis and cell growth: mTOR coordinates transcription by all three classes of nuclear RNA polymerases. Oncogene 25, 6384–6391 (2006). 10.1038/sj.onc.1209883

20 Liu, Y. et al. Single-cell transcriptome reveals insights into the development and function of the zebrafish ovary. Elife 11 (2022). 10.7554/eLife.76014

21 Bartsch D, K. K., Ahuja G, Lackmann JW, Hescheler J, Weber T, Bazzi H, Clamer M, Mendjan S, Papantonis A, Kurian L. mRNA translational specialization by RBPMS presets the competence for cardiac commitment in hESCs. Science Advances 9, 1–17 (2023). 10.1126/sciadv.ade1792

22 Farazi, T. A. et al. Identification of the RNA recognition element of the RBPMS family of RNA-binding proteins and their transcriptome-wide mRNA targets. Rna (2014). 10.1261/rna.045005.114

23 Heim, A. E. et al. Oocyte polarity requires a Bucky ball-dependent feedback amplification loop. *Development (Cambridge*, England) 141, 842–854 (2014).

24 Bertho, S. et al. A transgenic system for targeted ablation of reproductive and maternal-effect genes. Development 148 (2021). 10.1242/dev.198010

25 Jang, S. et al. The *Drosophila* ribosome protein S5 paralog RpS5b promotes germ cell and follicle cell differentiation during oogenesis. Development 148 (2021). 10.1242/dev.199511

26 Kong, J., et al. A ribosomal protein S5 isoform is essential for oogenesis and interacts with distinct RNAs in Drosophila melanogaster. Scientific Reports 9 (2019). 10.1038/s41598-019-50357-z

27 Deng M, W. X., Xiong Z, Tang P. Control of RNA degradation in cell fate decision. Front Cell Dev Biol (2023). 10.3389/fcell.2023.1164546

28 Ivanova, I. et al. The RNA m 6 A Reader YTHDF2 Is Essential for the Post-transcriptional Regulation of the Maternal Transcriptome and Oocyte Competence. Molecular Cell 67, 1059–1067.e1054 (2017). 10.1016/j.molcel.2017.08.003

29 Morgan, M. et al. mRNA 3ʹ uridylation and poly(A) tail length sculpt the mammalian maternal transcriptome. Nature 548, 347–351 (2017). 10.1038/nature23318

30 Blatt P, W.-D. S., McCarthy A, Breznak S, Hurton MD, Upadhyay M, Bennink B, Camacho J, Lee MT, Rangan P. RNA degradation is required for the germ-cell to maternal transition in Drosophila. Curr Biol. 31, 2984–2994 (2021). 10.1016/j.cub.2021.04.052

31 Iadevaia V, L. R., Proud CG. mTORC1 signaling controls multiple steps in ribosome biogenesis. Semin Cell Dev Biol 36, 113–120 (2014). 10.1016/j.semcdb.2014.08.004

32 Mehlmann, L. M. Stops and starts in mammalian oocytes: recent advances in understanding the regulation of meiotic arrest and oocyte maturation. Reproduction 130, 791–799 (2005). 10.1530/rep.1.00793

33 Lafontaine, D. L. J., Riback, J. A., Bascetin, R. & Brangwynne, C. P. The nucleolus as a multiphase liquid condensate. Nat Rev Mol Cell Biol 22, 165–182 (2021). 10.1038/s41580-020-0272-6

34 Raikova, E. V., Steinert, G. & Thomas, C. Amplified ribosomal DNA in meiotic prophase oocyte nuclei of acipenserid fishes. Wilehm Roux Arch Dev Biol 186, 81–85 (1979). 10.1007/BF00848111

35 Thiry, M. & Poncin, P. Morphological changes of the nucleolus during oogenesis in oviparous teleost fish, Barbus barbus (L.). J Struct Biol 152, 1–13 (2005). 10.1016/j.jsb.2005.07.006

36 Thiry, M. & Lafontaine, D. L. Birth of a nucleolus: the evolution of nucleolar compartments. Trends Cell Biol 15, 194–199 (2005). 10.1016/j.tcb.2005.02.007

37 Ochs, R. L., Lischwe, M. A., Spohn, W. H. & Busch, H. Fibrillarin: a new protein of the nucleolus identified by autoimmune sera. Biol Cell 54, 123–133 (1985). 10.1111/j.1768-322x.1985.tb00387.x

38 Iadevaia, V., Zhang, Z., Jan, E. & Proud, C. G. mTOR signaling regulates the processing of pre-rRNA in human cells. Nucleic Acids Res 40, 2527–2539 (2012). 10.1093/nar/gkr1040

39 Iida, T. & Lilly, M. A. missing oocyte encodes a highly conserved nuclear protein required for the maintenance of the meiotic cycle and oocyte identity in Drosophila. Development 131, 1029–1039 (2004). 10.1242/dev.01001

40 Kettleborough, R. N. et al. A systematic genome-wide analysis of zebrafish protein-coding gene function. Nature 496, 494–497 (2013). 10.1038/nature11992

41 Hwang, W. Y. et al. Heritable and precise zebrafish genome editing using a CRISPR-Cas system. PloS one 8, e68708 (2013). 10.1371/journal.pone.0068708

42 Leu, D. H. & Draper, B. W. The ziwi promoter drives germline-specific gene expression in zebrafish. Developmental dynamics : an official publication of the American Association of Anatomists 239, 2714–2721 (2010). 10.1002/dvdy.22404

43 Grabiner, B. C. et al. A diverse array of cancer-associated MTOR mutations are hyperactivating and can predict rapamycin sensitivity. Cancer Discov 4, 554–563 (2014). 10.1158/2159-8290.CD-13-0929

44 Bar-Peled, L. et al. A Tumor suppressor complex with GAP activity for the Rag GTPases that signal amino acid sufficiency to mTORC1. Science 340, 1100–1106 (2013). 10.1126/science.1232044

45 Wei, Y. et al. The GATOR complex regulates an essential response to meiotic double-stranded breaks in Drosophila. Elife 8 (2019). 10.7554/eLife.42149

46 McKim, K. S. & Hayashi-Hagihara, A. mei-W68 in Drosophila melanogaster encodes a Spo11 homolog: evidence that the mechanism for initiating meiotic recombination is conserved. Genes Dev 12, 2932–2942 (1998). 10.1101/gad.12.18.2932

47 Sekelsky, J. J. et al. Identification of novel Drosophila meiotic genes recovered in a P-element screen. Genetics 152, 529–542 (1999). 10.1093/genetics/152.2.529

48 Blokhina, Y. P., Nguyen, A. D., Draper, B. W. & Burgess, S. M. The telomere bouquet is a hub where meiotic double-strand breaks, synapsis, and stable homolog juxtaposition are coordinated in the zebrafish, Danio rerio. PLoS Genet 15, e1007730 (2019). 10.1371/journal.pgen.1007730

49 Kim, S. H., Speirs, C. K., Solnica-Krezel, L. & Ess, K. C. Zebrafish model of tuberous sclerosis complex reveals cell-autonomous and non-cell-autonomous functions of mutant tuberin. Dis Model Mech 4, 255–267 (2011). 10.1242/dmm.005587

50 Nie, D. et al. Tsc2-Rheb signaling regulates EphA-mediated axon guidance. Nat Neurosci 13, 163–172 (2010). 10.1038/nn.2477

51 Bashamboo, A., Eozenou, C., Rojo, S. & McElreavey, K. Anomalies in human sex determination provide unique insights into the complex genetic interactions of early gonad development. Clin Genet 91, 143–156 (2017). 10.1111/cge.12932

52 Sinclair, A. & Smith, C. Females battle to suppress their inner male. Cell 139, 1051–1053 (2009). 10.1016/j.cell.2009.11.036

53 Astapova, O., Minor, B. M. N. & Hammes, S. R. Physiological and Pathological Androgen Actions in the Ovary. Endocrinology 160, 1166–1174 (2019). 10.1210/en.2019-00101

54 Mercer, M., Jang, S., Ni, C. & Buszczak, M. The Dynamic Regulation of mRNA Translation and Ribosome Biogenesis During Germ Cell Development and Reproductive Aging. Front Cell Dev Biol 9, 710186 (2021). 10.3389/fcell.2021.710186

55 Falahati, H., Pelham-Webb, B., Blythe, S. & Wieschaus, E. Nucleation by rRNA Dictates the Precision of Nucleolus Assembly. Curr Biol 26, 277–285 (2016). 10.1016/j.cub.2015.11.065

56 Ortega-Recalde, O., Day, R. C., Gemmell, N. J. & Hore, T. A. Zebrafish preserve global germline DNA methylation while sex-linked rDNA is amplified and demethylated during feminisation. Nat Commun 10, 3053 (2019). 10.1038/s41467-019-10894-7

57 Xiong, M. et al. Conditional ablation of Raptor in the male germline causes infertility due to meiotic arrest and impaired inactivation of sex chromosomes. FASEB J 31, 3934–3949 (2017). 10.1096/fj.201700251R

58 Zhu, Z. et al. Rapamycin-mediated mTOR inhibition impairs silencing of sex chromosomes and the pachytene piRNA pathway in the mouse testis. Aging (Albany NY) 11, 185–208 (2019). 10.18632/aging.101740

59 Yang, Q. et al. Rapamycin improves the quality and developmental competence of in vitro matured oocytes in aged mice and humans. Aging (Albany NY) 14, 9200–9209 (2022). 10.18632/aging.204401

60 Westerfield, M. The Zebrafish Book. A Guide for the Laboratory Use of Zebrafish (Danio rerio), 4th Edition., (University of Oregon Press, 2000).

61 Kimmel, C. B., Ballard, W. W., Kimmel, S. R., Ullmann, B. & Schilling, T. F. Stages of embryonic development of the zebrafish. Developmental dynamics : an official publication of the American Association of Anatomists 203, 253–310 (1995). 10.1002/aja.1002030302

62 Gagnon, J. A. et al. Efficient mutagenesis by Cas9 protein-mediated oligonucleotide insertion and large-scale assessment of single-guide RNAs. PloS one 9, e98186 (2014). 10.1371/journal.pone.0098186

63 Montague, T. G., Cruz, J. M., Gagnon, J. A., Church, G. M. & Valen, E. CHOPCHOP: a CRISPR/Cas9 and TALEN web tool for genome editing. Nucleic Acids Res 42, W401–407 (2014). 10.1093/nar/gku410

64 Jinek, M. et al. A programmable dual-RNA-guided DNA endonuclease in adaptive bacterial immunity. Science 337, 816–821 (2012). 10.1126/science.1225829

65 Truett, G. E. et al. Preparation of PCR-quality mouse genomic DNA with hot sodium hydroxide and tris (HotSHOT). Biotechniques 29, 52, 54 (2000).

66 Song, H. W. et al. Hermes RNA-binding protein targets RNAs-encoding proteins involved in meiotic maturation, early cleavage, and germline development. Differentiation 75, 519–528 (2007). 10.1111/j.1432-0436.2006.00155.x

67 Wu, T. et al. clusterProfiler 4.0: A universal enrichment tool for interpreting omics data. Innovation (Camb) 2, 100141 (2021). 10.1016/j.xinn.2021.100141

68 Yu, G., Wang, L. G., Han, Y. & He, Q. Y. clusterProfiler: an R package for comparing biological themes among gene clusters. OMICS 16, 284–287 (2012). 10.1089/omi.2011.0118

69 Grote, S. GOfuncR: Gene ontology enrichment using FUNC. R package version 1.18.0., <10.18129/B9.bioc.GOfuncR> (2022).

70 Walter, W., Sanchez-Cabo, F. & Ricote, M. GOplot: an R package for visually combining expression data with functional analysis. Bioinformatics 31, 2912–2914 (2015). 10.1093/bioinformatics/btv300

71 Liu, Y. & Li, G. Empowering biologists to decode omics data: the Genekitr R package and web server. BMC Bioinformatics 24, 214 (2023). 10.1186/s12859-023-05342-9

72 Sayols, S. rrvgo: a Bioconductor package for interpreting lists of Gene Ontology terms. MicroPubl Biol 2023 (2023). 10.17912/micropub.biology.000811

73 K Blighe, S. R., M Lewis EnhancedVolcano: Publication-ready volcano plots with enhanced colouring and labeling. R package version 1.20.0, <10.18129/B9.bioc.EnhancedVolcano> (2023).

74 ggplot2 : Elegant Graphics for Data Analysis v. 2nd 2016. (Springer Springer International Publishing : Imprint: Springer, Cham, 2016).

75 Biodock. Biodock, AI Software Platform, <Available from www.biodock.ai> (2023).

