## Supplementary material for "Rbpms2 promotes female fate upstream of the nutrient sensing Gator2 complex component, Mios": Wilson et al Supplemental Materials

### Supplemental Figures and Materials

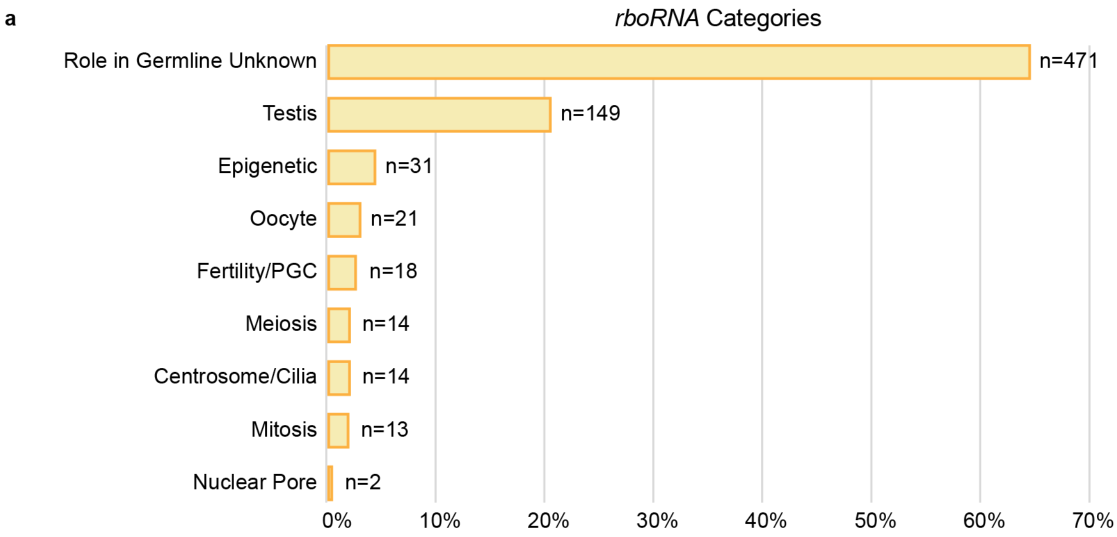


**Supplemental Figure 1. Several *rboRNAs* are associated with testis fates.** (a) Categorization of *rboRNAs* by literature search. Number of *rboRNAs* in each category are indicated.

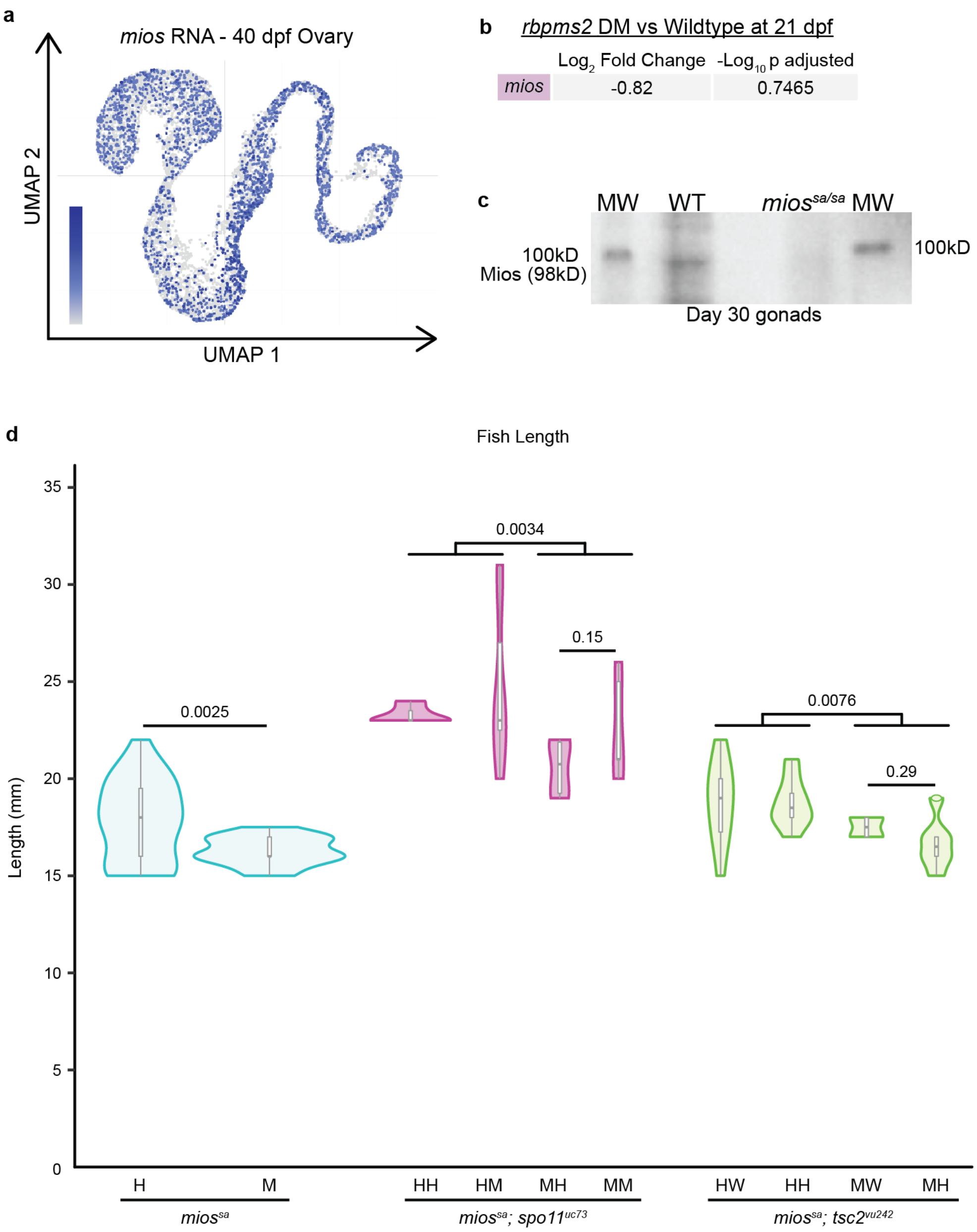


**Supplemental Figure 2. Mios is absent in *mios*$^{sa/sa}$ fish and loss of Mios reduces fish growth.** (a) UMAP of *mios* in the 40 dpf ovary[20]. (b) $Log_2$ Fold Change and $-Log_{10}$ p adjusted values of *mios* RNA expression from bulk RNA sequencing of *rbpms2* DMs versus wild-type gonads at 21 dpf. (c) Western blot of Mios in 30 dpf wild-type and *mios*$^{sa/sa}$ gonads. (d) Lengths of *mios*$^{sa}$, *mios*$^{sa}$*; spo11*$^{uc73}$, and *mios*$^{sa}$*; tsc2*$^{vu242}$ fish. A minimum of 4 fish per genotype were measured. Two-tailed paired equal variance student's t tests were performed for the indicated groups with a significant p-value of less than 0.05.

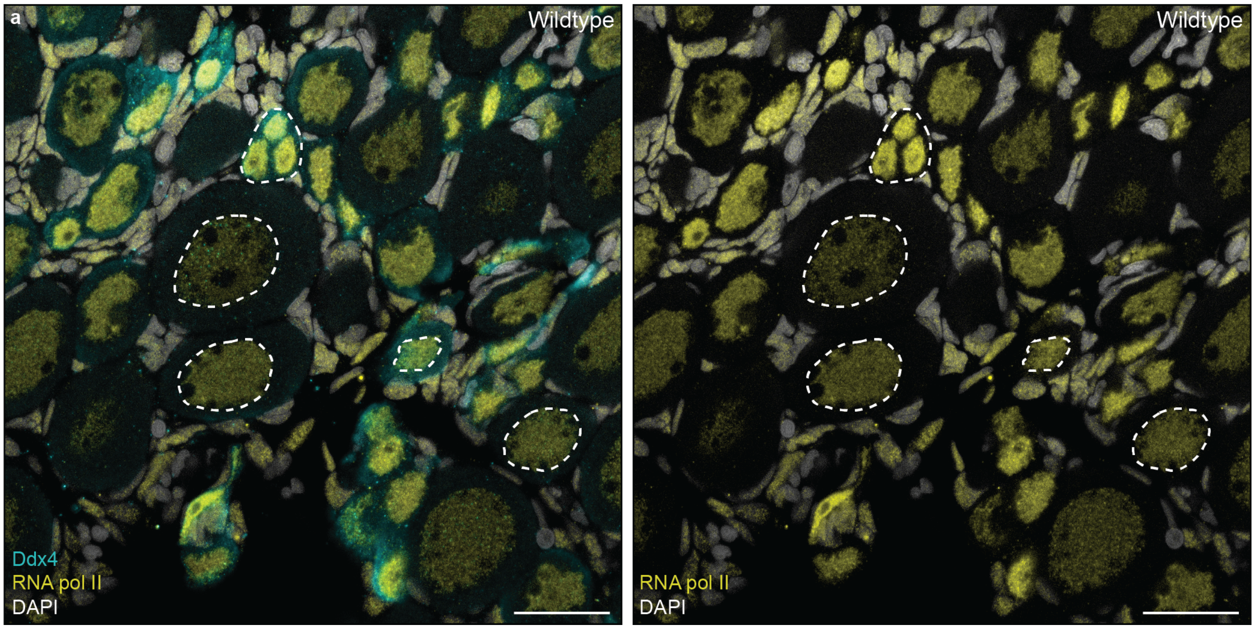


**Supplemental Figure 3. RNA pol II localization is highest in prophase I oocytes.** (a) RNA pol II (yellow) localization in 35 dpf wildtype gonads. Ddx4 labels germ cells (teal) and DAPI labels nuclei (white). Dashed white lines indicate nuclei of different cells stages and adjacent panel shows RNA pol II and DAPI localization in indicated cells. Scale bar is 50 μM.

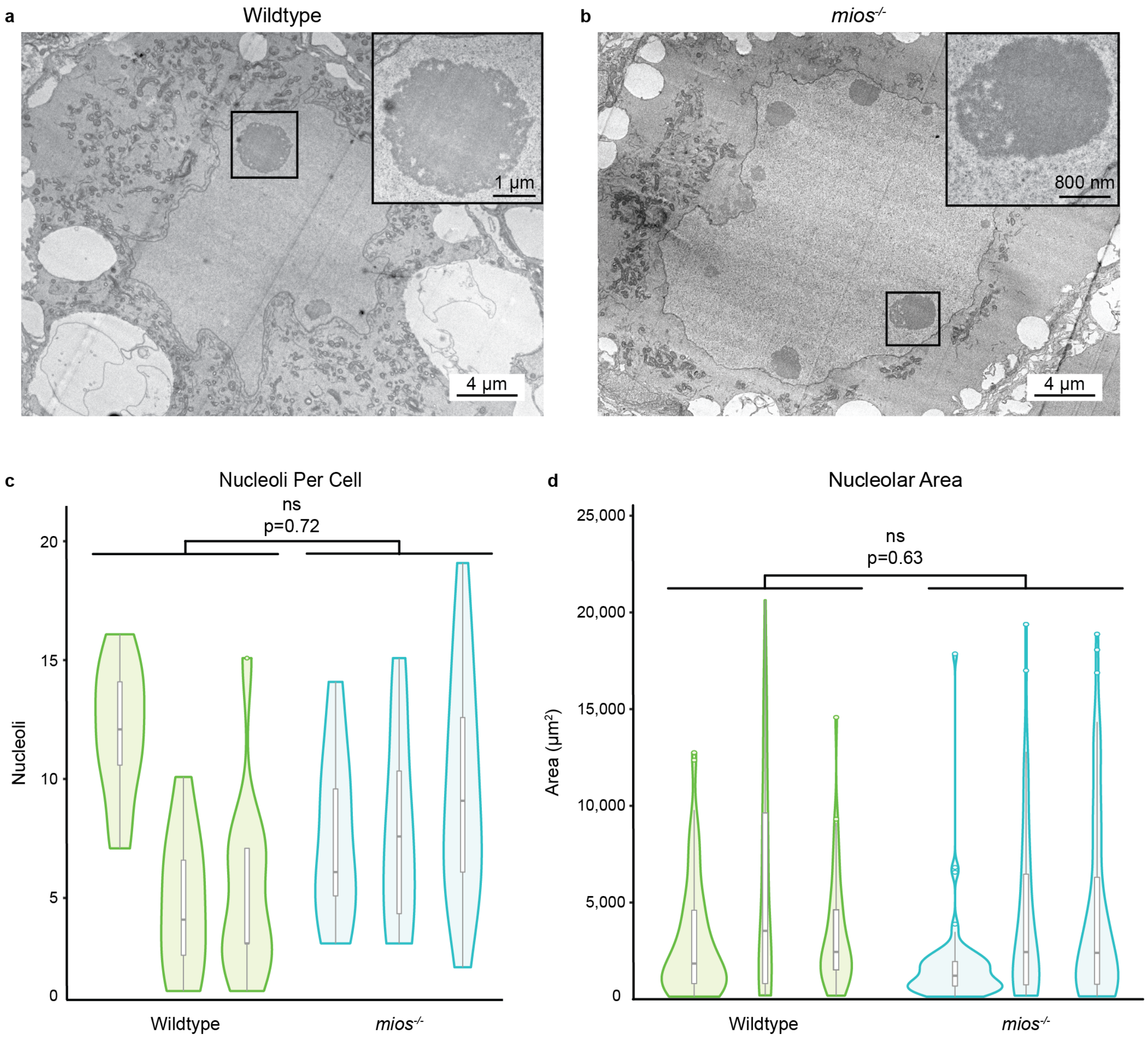


**Supplemental Figure 4. Nucleoli do not show significant ultrastructural differences between wildtype and *mios*$^{-/-}$ oocytes.** (a-b) Transmission electron micrographs of 35 dpf wildtype and *mios*$^{-/-}$ oocytes. Inset shows representative nucleolus for each genotype and corresponding scale bar. (c) Quantification of nucleoli per cell for 35 dpf wildtype (n=3) and *mios*$^{-/-}$ (n=3) oocytes. (d) Quantification of nucleolar area from 35 dpf wildtype (n=3) and *mios*$^{-/-}$ (n=3) oocytes. (c-d) Nucleoli from a minimum of 10 cells per fish were analyzed. Two tailed paired equal variance student's t-tests were performed for the indicated groups with a significant p-value of less than 0.05.

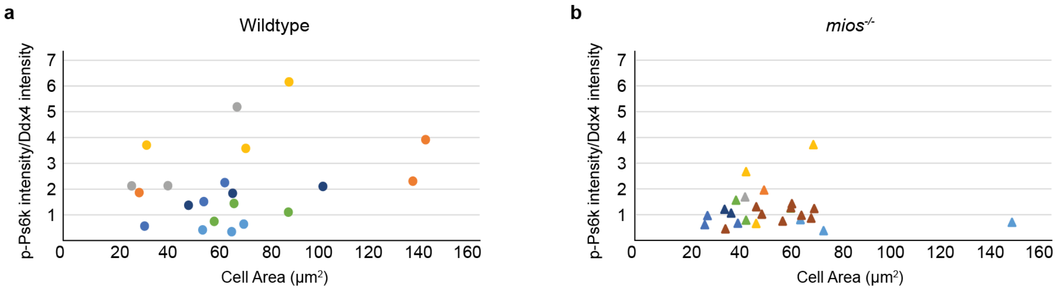


**Supplemental Figure 5. *mios*$^{-/-}$ oocytes have a significant decrease in p-Ps6k.** (a-b) distribution of p-Ps6k fluorescence intensity normalized to Ddx4 fluorescence intensity and correlated to (a) wild-type and (b) *mios*$^{-/-}$ oocyte size. (a) Circles indicate individual cells (n=21 total) and colors correspond to the specific fish (n=7 total). (b) Triangles indicate individual cells (n=25 total) and colors correspond to the specific fish (n=8 total).

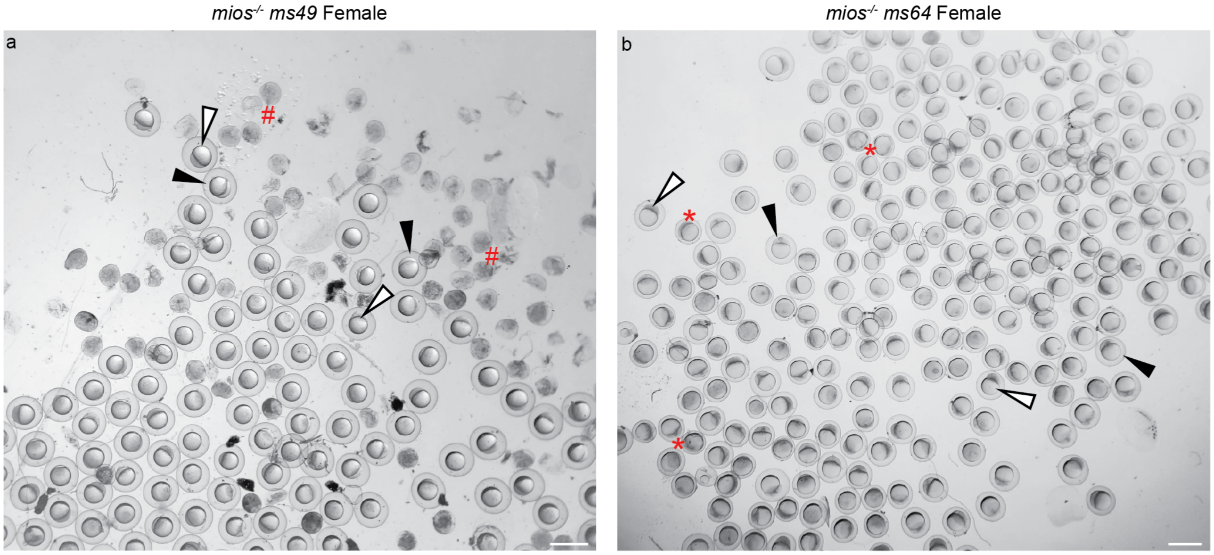


**Supplemental Figure 6. *mTOR*$^{ca}$ *ms49* and *ms64* *mios*$^{-/-}$ egg phenotypes.** (a) Representative eggs from a *mios*$^{-/-}$ *mTOR*$^{ca}$ *ms49* female. (b) representative eggs from a *mios*$^{-/-}$ *mTOR*$^{ca}$ *ms64* female. (a-b) White arrows with black outlines indicate activated, fertilized eggs, solid black arrows indicate activated, unfertilized eggs, (a) # indicates degenerating eggs, and (b) * indicates eggs with activation deficits. Scale bar is 1mm.

**Supplemental Table 1. Guide RNAs and Genotyping Primers.**

| Oligo Name | Sequence (5' to 3') |
|---|---|
| *mios sgRNA_1* | atttaggtgacactataGAGCTCAGTCTGTACCGGATgttttagagctagaaatagcaag |
| *mios sgRNA_2* | atttaggtgacactataAGGCCACGCATTTCATAAAGgttttagagctagaaatagcaag |
| *mios sgRNA_1 and 2_F* | CCAGATATCCTGTGGTCTCCTC |
| *mios sgRNA_1_R1* | CCACAGCTAACAAACACTCTGG |
| *mios sgRNA_1_R2* | CACAGCTAACAAACACTCTGGC |
| *mios sgRNA_2_R1* | CCTTACTTTTGGAGTTGTGGCT |
| *mios sgRNA_2_R2* | TGTCCCACAGCTAACAAACACT |
| *mios cDNA_F* | ACAAGCCACTGTCCTGCC |
| *mios cDNA_R* | TACGCTATCTGAAGGCACCAG |
| *rbpms2a ae30 F* | TTTGCTAAAGCCAACACGAA |
| *rbpms2a ae30 R* | ATTCACCCTGGCCAGAGTTT |
| *rbpms2b sa F_Mutant Assay* | CACTTATCAAGCTAACTTCAAAGCAGA |
| *rbpms2b sa F_Wildtype Assay* | TTTCAGGGTTATGAGGGTTCA |
| *rbpms2b sa R* | GCCAAAAGCAAAATATCAAACA |
| *mios ms20 F* | CCAGATATCCTGTGGTCTCCTC |
| *mios ms20 R* | CATTGGTATGTCCCACAGCTAA |
| *mios sa F* | GCCGAGTCGTGTTAACCACTT |
| *mios sa R1* | GCCAGCCAGTTACTGTCCAC |
| *mios sa R2* | ACGTCTTCTGGCTGGTGTTA |
| *spo11 uc73 F* | TCACAGCCAGGATGTTTTGA |
| *spo11 uc73 R* | CACCTGACATTGTTCCAGCA |
| *tsc2 vu242 F* | CCAGCACCACCTGCAGTCTGG |
| *tsc2 vu242 R* | CTCTTGGGCAGAGCAGAGAAGTTGG |
